# An immune-lncRNA risk model to predict prognosis for patients with head and neck squamous cell carcinoma

**DOI:** 10.1101/2022.03.10.483771

**Authors:** Xu Jiang, Ying Qin, Wei Xia, Jian Li, Nan Shi, Qingyuan Feng, Anzhou Tang, Xiang Yi

**Author notes:** These authors contributed equally to this work (Xu Jiang, Ying Qin). Corresponding author (YX).

## Abstract

**Background:** long noncoding RNA (lncRNA) are closely correlated with the occurrence and development of tumors. The purpose of this research is to construct a risk model of immune-related lncRNAs that can be used for predicting the survival and prognosis of head and neck squamous cell carcinoma (HNSCC) patients and further exploring the immune function of immune-lncRNAs.

**Methods:** obtained detailed transcriptional data and clinical information for 545 cases of HNSCC from The Cancer Genome Atlas (TCGA). Combined lncRNAs and the immune gene sets from the Molecular Signatures Database (MSigDB) were employed to construct co□expression networks, the significant correlation of immune-lncRNAs obtained from the results of Cox regression analysis were used to build the model. Survival analysis and QRT-PCR were used to confirm the expression of immune-lncRNAs and optimize the risk model. Then single-cell RNA Sequencing (scRNA-seq) dataset analysis of laryngeal squamous cell carcinoma (LSCC), co-expression analysis, correlation analysis and immunohistochemistry were used to explore the immune function of immune-lncRNAs.

**Results:** A total of 13 immune-lncRNAs including AC104083.1、AC004148.2、AL357033.4、AC116914.2、AC024075.1、AC008115.3、PCED1B-AS1、RAB11B-AS1、AC004687.1、 AL450992.2、EP300-AS1、AC024075.2、AC136475.2 were identified to construct the risk model. In this model, low-risk patients had higher overall survival (OS, P=1.923e-12, hazard ratio=2.104, 95% CI: 1.622-2.730). AC004687.1 and PCED1B-AS1 were overexpressed in T cells and were closely related to the genes encoding major histocompatibility complex II (MHC II) molecules. And the results of QRT-PCR and immunohistochemistry showed the expression significant positively correlated with HLA-DR which is one of the MHC II classical molecules.

**Conclusion:** the immune-associated risk model constructed by 13 lncRNAs has good prognostic value for HNSCC. AC004687.1 and PCED1B-AS1 are significant correlating to clinical characteristics and MHC II, we provide a new insight into the immune function of lncRNAs.

## INTRODUCTION

Head and neck squamous-cell carcinoma (HNSCC) is the sixth most common type of cancer worldwide and influence 600,000 new patients every year(1). HNSCC arises in the oral cavity, oropharynx, hypopharynx and larynx with poor prognosis, and the five-year survival rate is less than 50%(2). The main therapy for HNSCC currently combines surgical excision, radiation therapy, chemotherapy and biotherapy(3). In recent years, immunotherapy for HNSCC has attracted considerable attention, and the most widely used immunotherapy for cancer is moAb therapy(4). Cetuximab is the main immunotherapy medicine for HNSCC(5). EGFR, as an attractive target, is overexpressed in 80∼90% HNSCC(6), which causes proliferation and invasion of tumor cells and leads to poor survival and prognosis(7). Immunotherapy has had a positive effect on patients with HNSCC. Inhibitors of immune checkpoints, such as anti-CTLA-4 and PD-1 monoclonal antibodies, regulate T cell function, thereby eliminating immunosuppression in the tumor microenvironment to increase the overall survival time and disease-free survival of patients(8).

Long noncoding RNAs (lncRNAs) are a class of functional RNA molecules(9) that have limited protein coding ability and play dual roles in protein coding and noncoding(10). However, the functional representation of most lncRNAs has not been achieved to date. Some lncRNAs are associated with a series of biological processes, such as chromatin modification, transcription factor regulation and cell signal transduction(11). Increasing research suggests that a number of lncRNAs could be used as an important driving force of malignant transformation in cancer. The expression of some lncRNAs is associated with the characteristics of tumor growth and the survival rate of patients, which means that these lncRNAs will be used as a convenient biomarker of prognosis(12). For head and neck squamous-cell carcinoma, research suggests that the lncRNA HOTTIP is related to the progression and prognosis of tongue squamous-cell carcinoma(13). Some lncRNAs were certified to be abnormally expressed in oral cancer and metastatic cases, and lncRNAs in saliva are used as potential biomarkers for the diagnosis of oral squamous-cell carcinoma(14).

The transcription of some lincRNAs has been reported to participate in the immune response. For example, lincRNA-Cox2 has been identified as the key component of the inflammatory response(15). LincRNA-EPS inhibition is an inflammatory response inhibitor(16). Lnc-Lsm3b can compete with viral RNAs in the binding of RIG-I monomers and lead to RIG-I congenital function inactivation at the late stage of feedback in the innate response, which revealed noncanonical self-recognition in immunoregulation(17). NKILA can modulate the sensitivity of T cells to AICD by inhibiting the activity of NF-κB(18) and combine with NF-κB/IκB, directly covering the phosphorylation motif of IκB and inhibiting the metastasis of breast cancer(19). Mirt2 can be used as the checkpoint to prevent abnormal activation of inflammation, as it is the potential regulatory factor of macrophage polarization, inhibit the activation of the NF-κB and MAPK pathways and limit the production of proinflammatory cytokines(20). Therefore, the abnormal expression of immune-lncRNAs may have prognostic value for HNSCC patients and enable them to be used as potential targets for immunotherapy. In previous reports, immune-associated lncRNA has been used to construct a risk model to predict tumor risk(21, 22).

In this research, we used transcriptional data and clinical information of HNSCC in The Cancer Genome Atlas (TCGA) to construct and verify the immune-lncRNA risk model for the survival and prognosis of HNSCC patients.

## MATERIALS AND METHODS

### Patients and datasets

Detailed transcriptional data and clinical information in 545 cases of head and Heck squamous-cell carcinoma were obtained from The Cancer Genome Atlas (TCGA)(23)(https://portal.gdc.cancer.gov/). Patients without survival data or ≤ 30 days were excluded; finally, a total of 490 patients were included in the analysis.

### Patients and tissue samples

HNSCC tissues and adjacent normal tissues were obtained from 17 patients who underwent HNSCC resection at otorhinolaryngology head and neck surgery in the First Affiliated Hospital of Guangxi Medical University during 2020.6 – 2020.10. Informed consent was obtained from each patient and the Medical Ethics Committee of the First Affiliated Hospital of Guangxi Medical University approved the study. All the subjects have provided appropriate informed consent. The clinicopathological information of 17 HNSCC patients available at Table S1. All patients had primary HNSCC and did not undergo preoperative chemotherapy and/or radiotherapy. The tissues and corresponding adjacent normal tissues were divided into two parallel parts: one part were formalin fixed and paraffin-embedded while the other part were frozen and stored at −□80 °C to extract genomic RNA.

### Immune-lncRNA identification and extraction

The transcriptome gene matrix was differentiated into a lncRNA expression matrix and a protein coding matrix. The expression of immune-related genes from the protein coding matrix was extracted by coexpression correlation analysis between the lncRNA expression matrix and the immune gene sets M19817 and M13664 from the Molecular Signatures Database (MSigDB) of GSEA (https://www.gsea-msigdb.org/gsea/msigdb/index.jsp) (MsigDB annotated by GSEA, the gene sets according to different functions were differentiated into eight gene sets that included immune-related genes(24). Immune-related lncRNAs were filtered out, and expression values were extracted according to cor>0.4 (P<0.001) to obtain an immune-lncRNA expression matrix.

### Construction of the prognosis-related lncRNA model

The survival status and survival time of immune-lncRNAs were analyzed by Cox univariate analysis. Filtering by P<0.01 and extraction of the expression of single-factor significant lncRNAs was undertaken. The coefficient of single factor significance lncRNAs was calculated by logarithmic conversion hazard ratio (HR) obtained from univariate Cox regression analysis. According to the Cox model AIC (Akaike information criterion) optima, the immune-lncRNAs that were used to build the model were identified. Based on the previously reported risk score formula(25-27), Risk score = βgene1 × exprgene1 + βgene2 × exprgene2 + · + βgenen × exprgenen. The risk score of patients was calculated (βgene is the coefficient of lncRNAs, and exprgene is the expression quantity of lncRNAs). According to the median of the risk score, patients were divided into a high-risk group and a low-risk group.

### Single-cell RNA-seq analysis

Single-cell transcriptome files of HNSCC tumors (GSE150321) were downloaded from the Gene Expression Omnibus (GEO) database (https://www.ncbi.nlm.nih.gov/geo/query/acc.cgi?acc=GSE150321)(28). And the data of 3,885 single cells from a patient with laryngeal squamous cell carcinoma (LSCC). The Seurat package was used to perform single-cell RNA-seq analysis. The cells were filtered according to the expression gene number > 50, less than 20% of the gene from the mitochondrial genome. To explore the distribution of lncRNA in different cell types.

### Function prediction of single immune-associated lncRNA

Pearson correlation coefficient was used to detect the expression level correlation between of the protein coding gene (PCGs) and immune-lncRNA which used to construct the risk model. |Pearson correlation coefficient| > 0.40 and P < 0.01 will considered to be the protein coding gene associated with the lncRNA. Gene ontology(GO)(29) and Kyoto Encyclopedia of Genes and Genomes (KEGG)(30) analysis used to analyze the function and pathway of a single lncRNA.

### RNA isolation and QRT-PCR

Total RNA from tissues was extracted with RNAsimple Total RNA Kit (TIANGEN BIOTECH, BEIJING, China) according to the manufacturer’s protocol. Reverse-transcribed lncRNA cDNA was obtained using lnRcute lncRNA First-Strand cDNA kit (TIANGEN BIOTECH, BEIJING, China). Quantitative PCR was performed by using the lnRcute lncRNA QRT-PCR Kit (SYBR Green)(TIANGEN BIOTECH, BEIJING, China)and Bio-Rad CFX Connect Real-Time PCR Detection System (Bio-Rad, USA). The expression of HLA-DRA and HLA-DRB1 which are the downstream gene of immune-lncRNAs in HNSCC and adjacent tissues were obtained using Talent QRT-PCR PreMix(SYBR Green) (TIANGEN BIOTECH, BEIJING, China). The expression of HLA-DRA, HLA-DRB1 and lncRNA were normalized against GAPDH. The experiment was repeated 3 times. The differences between groups were calculated with the comparative Ct method (2^−^ΔΔCT). Each experiment was performed in triplicate. All experiments were performed in triplicate. Primer sequences are shown in supplementary Table S1.

### Immunohistochemical staining

The tissues were dewaxed and stained with hematoxylin and eosin (HE). HLA-DR protein expression was determined by immunostaining using the rabbit anti-human HLA-DR (1: 500 dilution; EPR3692, ab92511, Abcam). The HRP-conjugated secondary antibody was visualized by development with diaminobenzidine (DAB; DAB-0031/1031, MXB, China). Image Pro Plus 6.0 software was used to analyze the tissue photos. Mean optical density (OD)(31) was used to determine the difference of HLA-DR protein expression between HNSCC tissues and adjacent normal tissues.

### Statistical analysis

Hazard ratio (HR) and 95% confidence interval (CI) obtained from univariate and multivariate Cox regression analysis(32, 33) The accuracy of the risk score for survival prediction was evaluated by Kaplan–Meier survival curve and ROC curve(34), and the acreage under the ROC curve (AUC) was calculated. Principal component analysis (PCA). The Seurat package, gene ontology (GO) and Kyoto Encyclopedia of Genes and Genomes (KEGG) analysis. The above analysis was performed by R software (version 3.6.2). Gene set enrichment analysis (GSEA) (http://software.broadinstitute.org/gsea/index.jsp) was used for functional annotation of high-low risk groups, and a P value <0.05 was considered to be significant. Statistical analysis of the experiment was performed Graph Pad Prism, version 8.0.2 (Graph Pad Software, San Diego, CA, USA) and SPSS 23.0. Use The ROUT method in Graph Pad Prism to remove outliers. Shapiro-Wilk normality test determines the normality of the cleaned data, The normal distribution uses paired t-test, and the non-normal distribution uses Wilcoxon matched-pairs signed rank test, and a P value < 0.05 was considered to be significant.

## RESULTS

### Preliminary construction of a prognostic risk model for immune-associated lncRNA

#### Coexpression networks of immune-lncRNAs

A total of 7009 lncRNAs were extracted from the transcriptional data of 490 patients. Immune-related gene sets IMMUNE_RESPONSE (M19817) and IMMUNE_SYSTEM_PROCESS (M13664) were obtained from the Molecular Signatures Database v7.0 of GSEA (https://www.gsea-msigdb.org/gsea/msigdb/index.jsp). A total of 331 immune-related genes and the expression data were extracted from the protein coding matrix and analyzed by coexpression correlation with the lncRNA expression matrix, and filtering was undertaken according to (cor>0.4, P<0.001). Finally, 341 immune-lncRNAs were identified. We identified 46 single factor significant immune-lncRNAs by univariate Cox analysis between immune-related lncRNAs with survival time and survival status, and filtering was performed according to P<0.01 (S1 Fig).

#### Identification of prognosis-related immune-lncRNAs to constructed risk model

According to the Cox model AIC (Akaike information criterion) optima, 20 immune-lncRNAs from 341 immune-related lncRNAs were identified to construct a risk model. The 17 immune-lncRNAs including high-risk lncRNA IER3-AS1, and low-risk lncRNAs AC104083.1, AL109811.3, AC004148.2, AC015849.3, AL357033.4, RAD51-AS1, AC116914.2, AC024075.1, AC008115.3, PCED1B-AS1, RAB11B-AS1, AC004687.1, AL450992.2, EP300-AS1, AC024075.2 and AC136475.2. According to the median risk score, 490 patients were divided into a high-risk group and a low-risk group. Survival analysis was performed in both groups. The results showed that the five-year survival rate of the low-risk group was higher than that of the high-risk group (P=5.33e-13, hazard ratio=1.653, 95% CI: 1.460-1.871, Fig1).

**Fig1.**
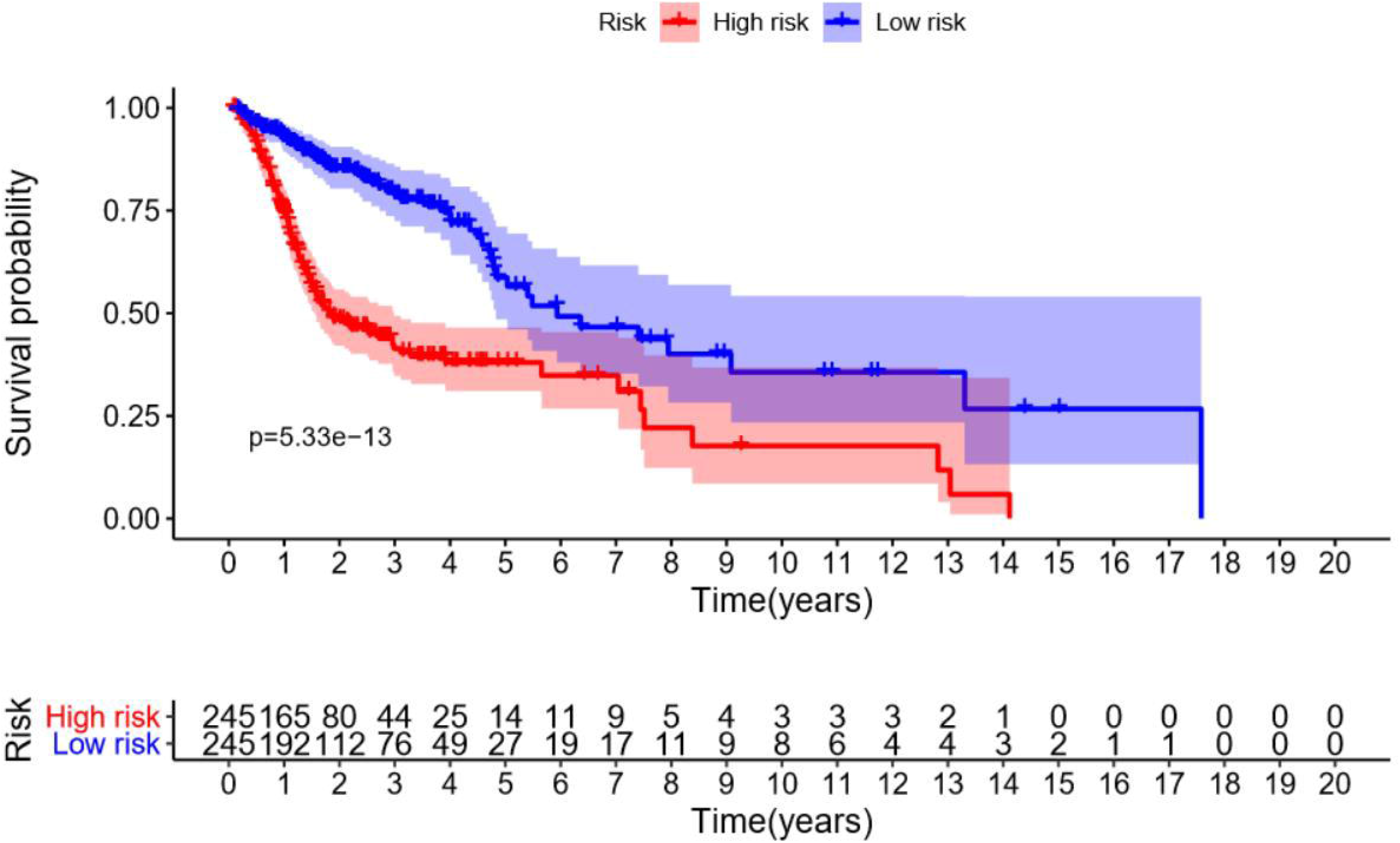
Survival curve of the high-risk group and low-risk group. The high-risk group showed a lower five-year survival rate than the low-risk group. The five-year survival rate of the high-risk group was approximately 34.8% (95% CI: 0.2674-0.454), and that of the low-risk group was approximately 56.5% (95% CI: 0.460-0.694), P=5.33e-13.

#### Analysis of independent prognostic

We combined the clinical characteristics and risk scores analyzed by independent prognosis with survival time and survival status. The clinical characteristics included age, gender, stage, T, N and extracapsular spread (ECS). M was excluded from the following analysis because only one patient had distant metastases. The results of univariate and multivariate factor independent prognostic analysis showed that the hazard ratio value of the risk score was 1.653 and 1.550 (P=1.94e-15, P=8.14e-11, Fig2A-B). These results indicate that the risk score of the model constructed by 17 immune-lncRNAs is a high-risk factor. The risk score can be used as an independent prognostic factor.

**Fig-2.**
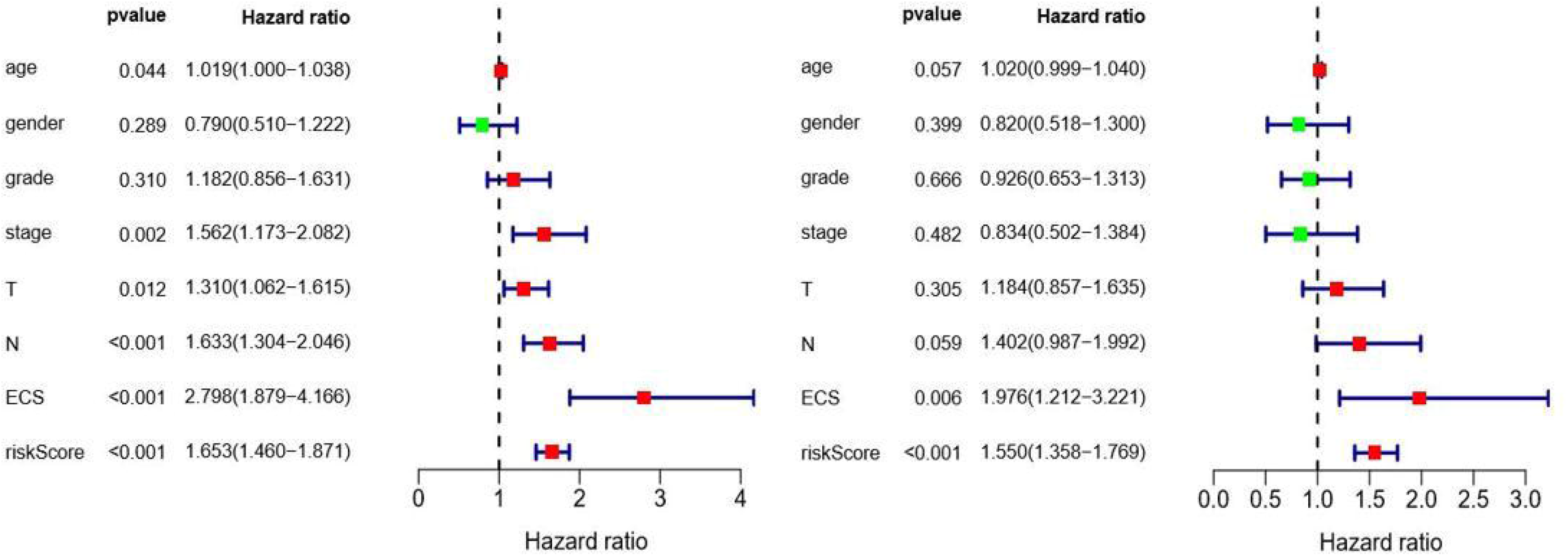
The results of independent prognostic Analysis. **(A)**Forest map of univariate factor independent prognostic analysis. **(B)**Forest map of multivariate factor independent prognostic analysis. A hazard ratio>1 indicates a high-risk factor. A hazard ratio <1 indicates a low-risk factor.

#### Correlation analysis of clinical characteristics

We assessed the correlation between 17 immune-lncRNAs and stage, T, N and ECS (Fig 3). The results showed that with increasing stage, the expression of AL357033.4 decreased (P=0.03, Fig 3A). As T increased, the expression of AC004687.1 (P=3.6e−05), AC024075.2(P=0.030), EP300-AS1(P=0.011) and PCED1B-AS1(P=0.024) decreased (Fig 3B). With increasing N, the expression of AL357033.4 decreased (P=0.006, Fig 3C). In lncRNAs with P<0.05, the expression of IER3-AS1(P=0.047) was higher in the patients with ECS, but the expression of AC024075.1(P=0.037) and AL357033.4(P=0.011) was lower in the patients without ECS (Fig 3D).

**Fig 3.**
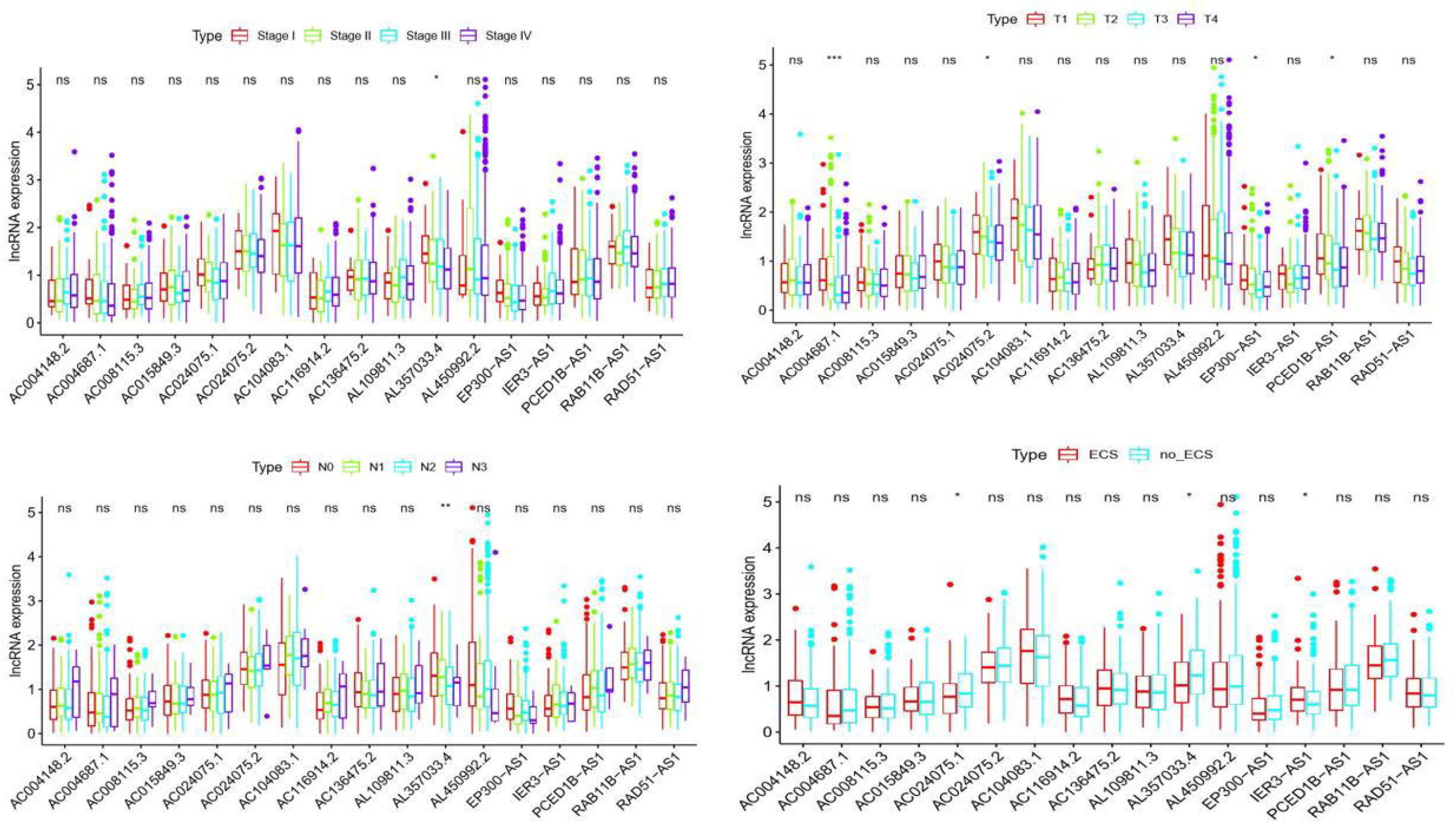
Results of the correlation analysis between the clinical characteristics of the 17 immune-lncRNAs. **(A)**Analysis with Stage. **(B)**Analysis with T. **(C)**Analysis with N. **(D)**Analysis with ECS. (ns:P>0.05, *: P<0.05, **: P<0.01, ***: P<0.001.)

### Optimization of immune-lncRNAs risk model

#### Survival analysis of single immune-lncRNA

Survival analysis was performed on the 17 immune-lncRNAs which were used to construct the risk model. The results showed that the high-risk lncRNA IER3-AS1 (P=4.17e−03) has worse survival and prognosis in the high-expression group than in the low-expression group, the differences were statistically significant. The 16 protective immune-lncRNAs, except for RAD51-AS1 and AL109811.3 had no significant effect on survival and prognosis, the other 14 including AC104083.1(P=2.408e−03), AC004148.2(P=9.19E-04), AC015849.3(P=7.557e−03), AL357033.4(P=1.42e−03), AC116914.2(P=1.101e−03), AC024075.1(P=3.244e−02), AC008115.3(P=1.646e−02), PCED1B-AS1(P=2.224e−02), RAB11B-AS1(P=2.782e−02), AC004687.1(P=2.767e−03), AL450992.2(P=3.355e−04), EP300-AS1(P=1.276e−02), AC024075. 2(P=3.302e−04) and AC136475.2(P=3.942e−03) have better survival and prognosis in the high-expression group than in the low-expression group, the differences were statistically significant. (S2 Fig)

#### Expression of survival-related lncRNAs in HNSCC specimens

Since most of these lncRNAs are unannotated, their expression in tissues was determined by QRT-PCR. Detected the 15 lncRNAs in survival analysis with statistically significant in 17 pairs of HNSCC cancer and adjacent normal tissues whether there were differences in expression. The results showed that the expression of AL357033.4(P=0.0016), AC024075.2(P=0.0024), AC004687.1(P=0.0174), IER3-AS1(P=0.0093), AC024075.1(P=0.0154), EP300-AS1(P=0.0067), AL450992.2(P=0.0323), AC008115.3(P=0.0001), AC004148.2(P=0.0073), RAB11B-AS1(P=0.0210), AC136475.2(P=0.0009), AC116914.2(P=0.0265), PCED1B-AS1(P=0.0032) and AC104083.1(P=0.0009) in normal tissue was higher than tumor tissue, the differences were statistically significant (Fig 4). All of them, except IER3-AS1, were consistent with the previous analysis that these lncRNA were protective lncRNA and had good survival and prognosis.

**Fig 4.**
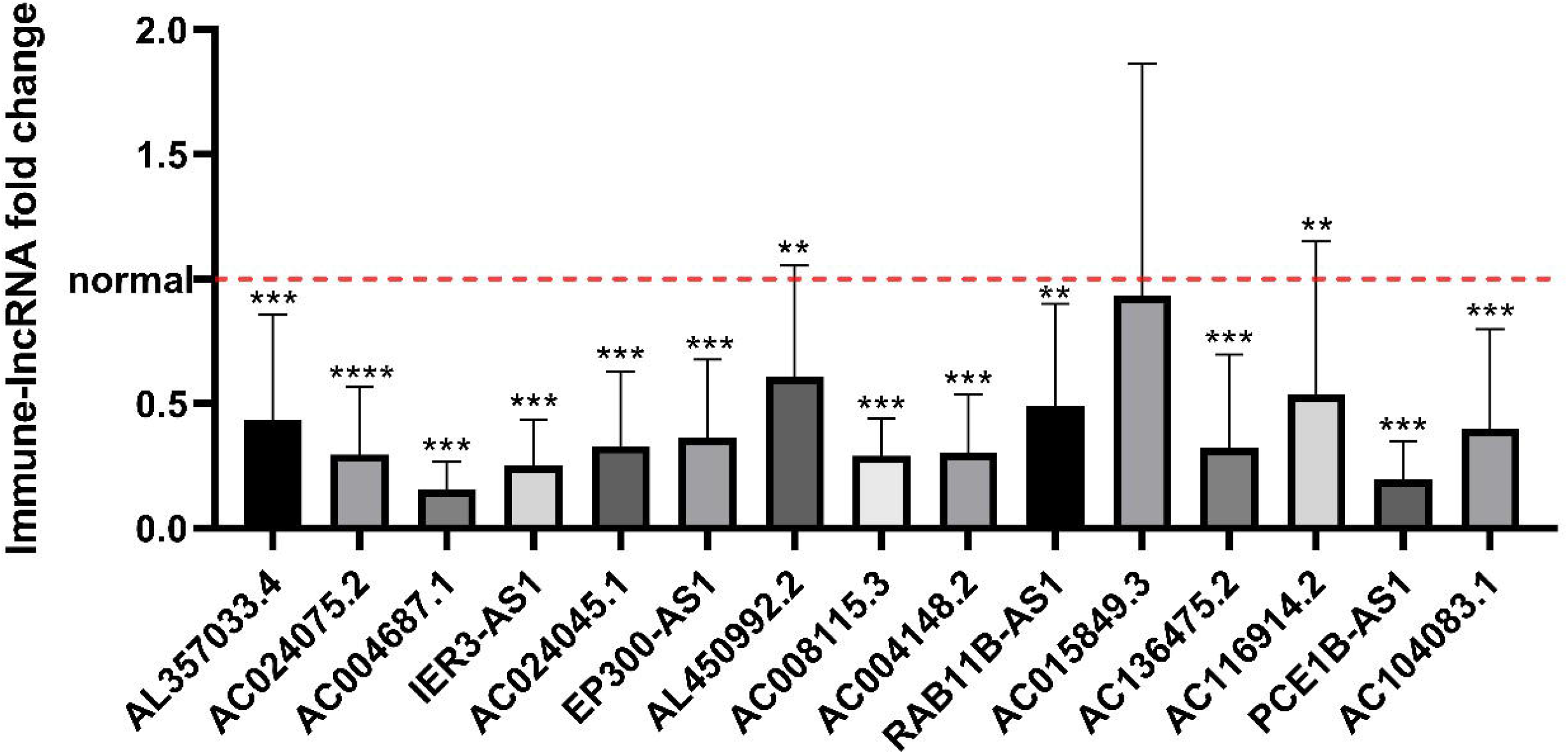
The expression of single lncRNA in tumor tissues compared with adjacent normal tissues was determined by QRT-PCR. *: P<0.05, **: P<0.01, ***:P <0.001, ****:P <0.0001.

#### Establish the immune-lncRNAs risk model

We reconstructed the risk model by immune-lncRNAs which has survival and prognosis significance and with differential expression in HNSCC cancer and adjacent normal tissues. IER3-AS1 as high-risk lncRNA but has less expressed in tumors than normal tissue and therefore is not included. Finally, 13 protective immune-lncRNAs included AC104083.1, AC004148.2, AL357033.4, AC116914.2, AC024075.1, AC008115.3, PCED1B-AS1, RAB11B-AS1, AC004687.1, AL45092.2, EP300-AS1, AC024075.2 and AC136475.2 used to construct the risk model. Survival analysis of 490 patients again showed the low-risk group had a higher five-year survival rate than the high-risk group (P=1.923e-12, hazard ratio=2.104, 95% CI: 1.622-2.730, Fig 5).

**Fig 5.**
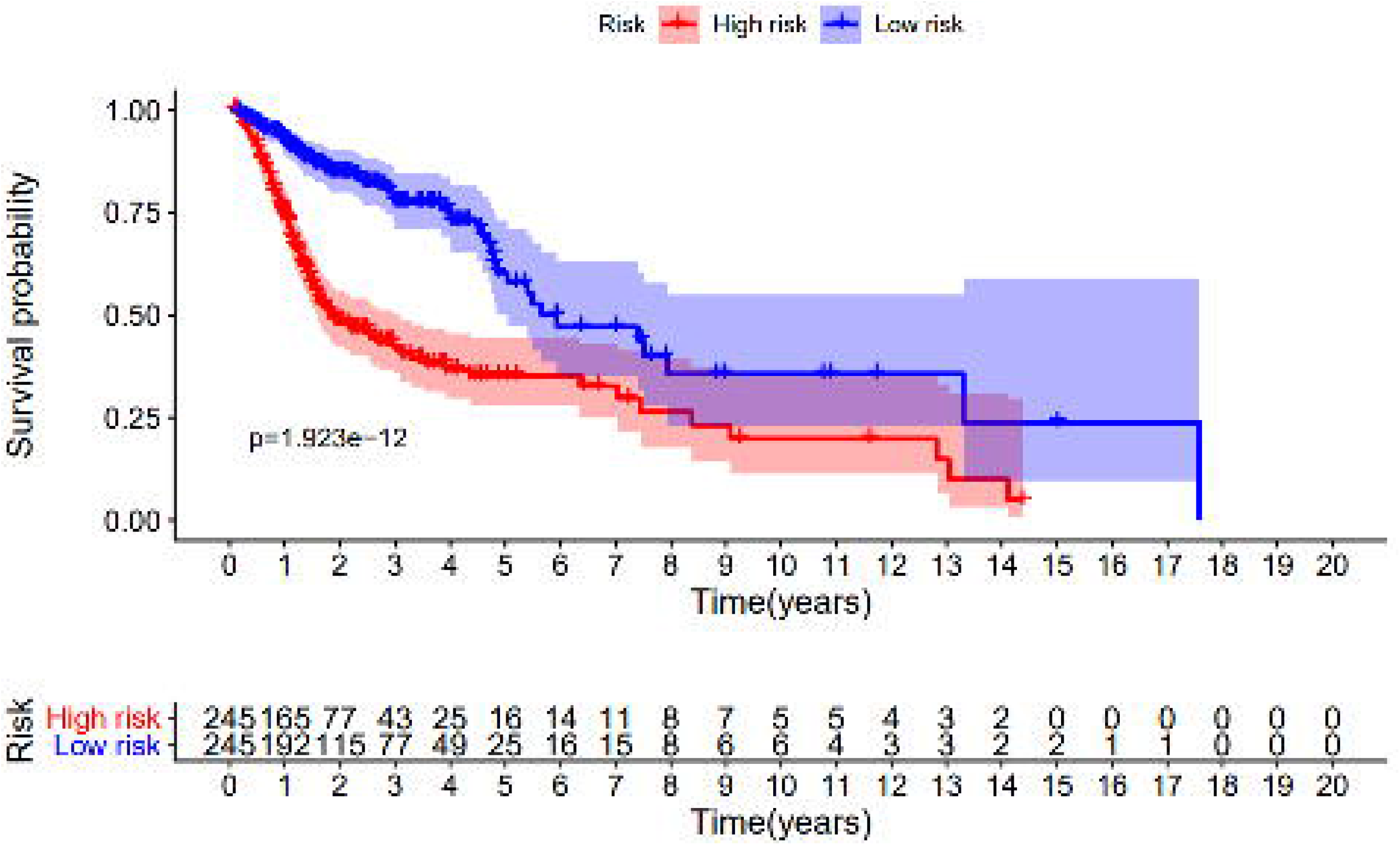
Survival curve of the high-risk group and low-risk group.

The high-risk group showed a lower five-year survival rate than the low-risk group. The five-year survival rate of the high-risk group was approximately 35.2% (95% CI: 0.279-0.443), and that of the low-risk group was approximately 71.1% (95% CI: 0.630-0.803), P=1.923e-12.

### Validation of immune-lncRNAs risk model

#### Analysis of independent prognostic

We combined the clinical characteristics and risk scores of optimized model analyzed by independent prognosis with survival time and survival status. The results of univariate and multivariate factor independent prognostic analysis showed that the hazard ratio value of the risk score was 2.104 and 2.072 (P=2.17e-08, P= 6.52e-07, Fig 6A-B), higher than before optimization. These results indicate that the risk score of the model constructed by 13 immune-lncRNAs is a high-risk factor. The risk score can be used as an independent prognostic factor. The risk score had the maximum area under the multiple index ROC curve (AUC=0.701). The results showed that the accuracy of the risk score was higher than that of other clinical characteristics in the prediction (Fig 6C).

**Fig 6.**
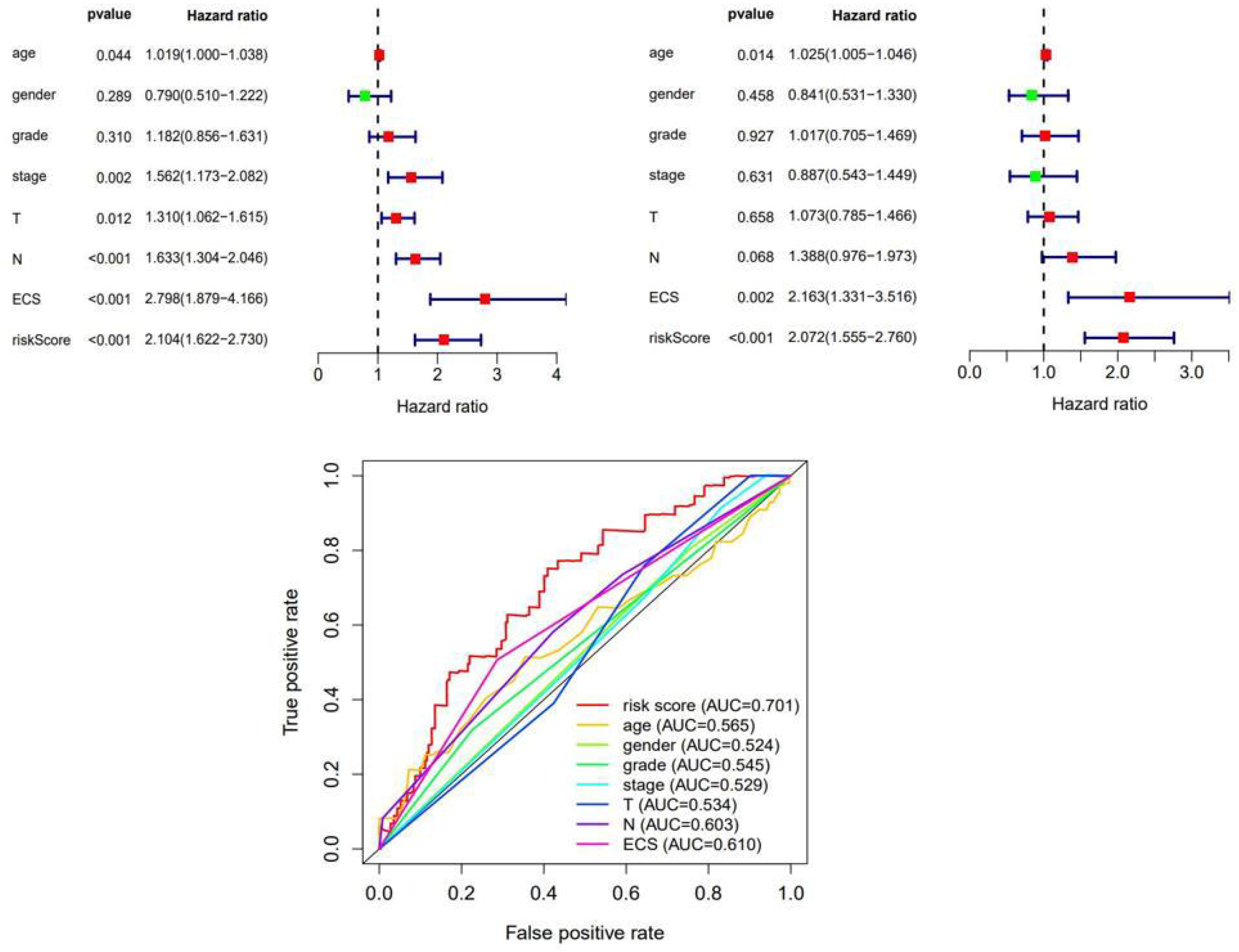
The results of independent prognostic Analysis. **(A)**Forest map of univariate factor independent prognostic analysis. **(B)**Forest map of multivariate factor independent prognostic analysis. A hazard ratio >1 indicates a high-risk factor. A hazard ratio <1 indicates a low-risk factor. **(C)**The multiple index ROC curve of true positive rate and false positive rate, risk score AUC=0.701.

#### Low□risk and high□risk groups displayed different immune statuses

We ranked patients from low to high risk and plotted a risk curve combining survival time, survival status and the expression data of 13 immune-lncRNAs (Fig 7A-B). The results showed that as the risk score of patients increases, the number of deaths increases. The expression data of 13 immune-lncRNAs showed differences between the low□risk and high□risk groups (Fig 7C).

**Fig 7.**
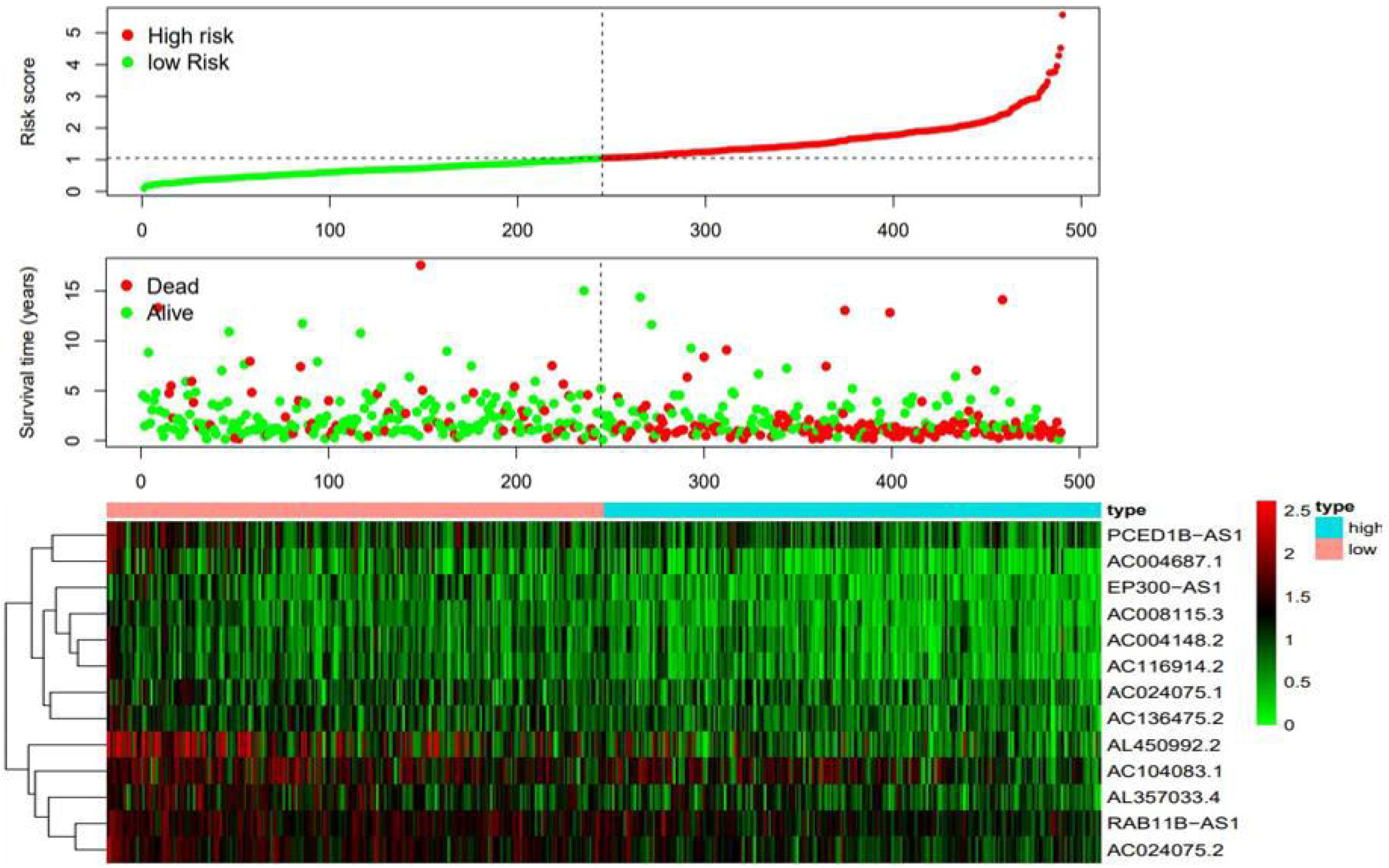
Risk curve. **(A)**Red indicates the high-risk group, green indicates the low-risk group, and the risk score increased from left to right. **(B)**The relationship between the survival status of patients and risk score. As the risk score increases from left to right, the number of deaths has increased, but the number of survivors has diminished. **(C)**Expression heat map of 13 immune-lncRNAs.

We conducted principal component analysis (PCA) to explore the difference between all the genes of 490 patients, immune-genes, immune-lncRNAs and the immune-lncRNAs used to construct the model and the risk score (Fig 8). The PCA results of the model constructed by 13 immune-lncRNAs showed that the low□risk and high□risk groups concentrated in different areas were more significant, which means that the model and risk score can be used to differentiate immune status between low□risk and high□risk groups of HNSCC patients more effectively.

**Fig 8.**
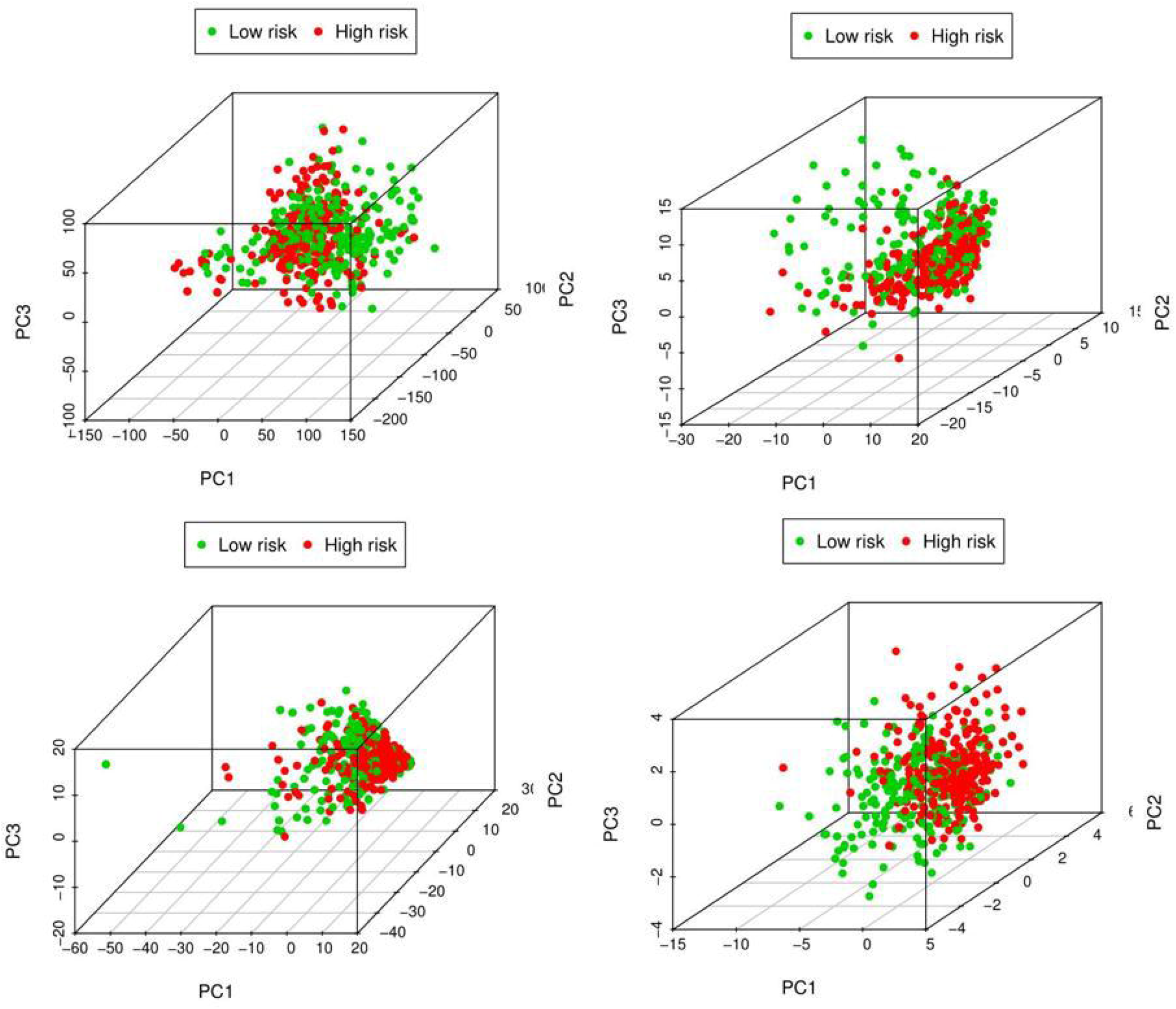
Results of PCA. **(A)**All the genes. **(B)**Immune genes. **(C)**Immune-lncRNAs. **(D)**Immune lncRNAs used to construct the model and risk score. Red indicates the high-risk group, and green indicates the low-risk group.

#### Functional annotation of GSEA

The immune-related gene sets IMMUNE_RESPONSE and IMMUNE_SYSTEM_PROCESS were used to perform functional annotation (S3 Fig). Although the p-value was not less than 0.05, in this risk model, the low-risk group showed more active trends in enrichment of immune response and immune system process than the high-risk group.

KEGG signaling pathway functional annotation was further performed by gene set enrichment analysis (GSEA). We focused on immune-related results (Fig 9A-B). The high□risk groups had no enrichment results. The results of the low□risk groups are the following:

**Fig 9.**
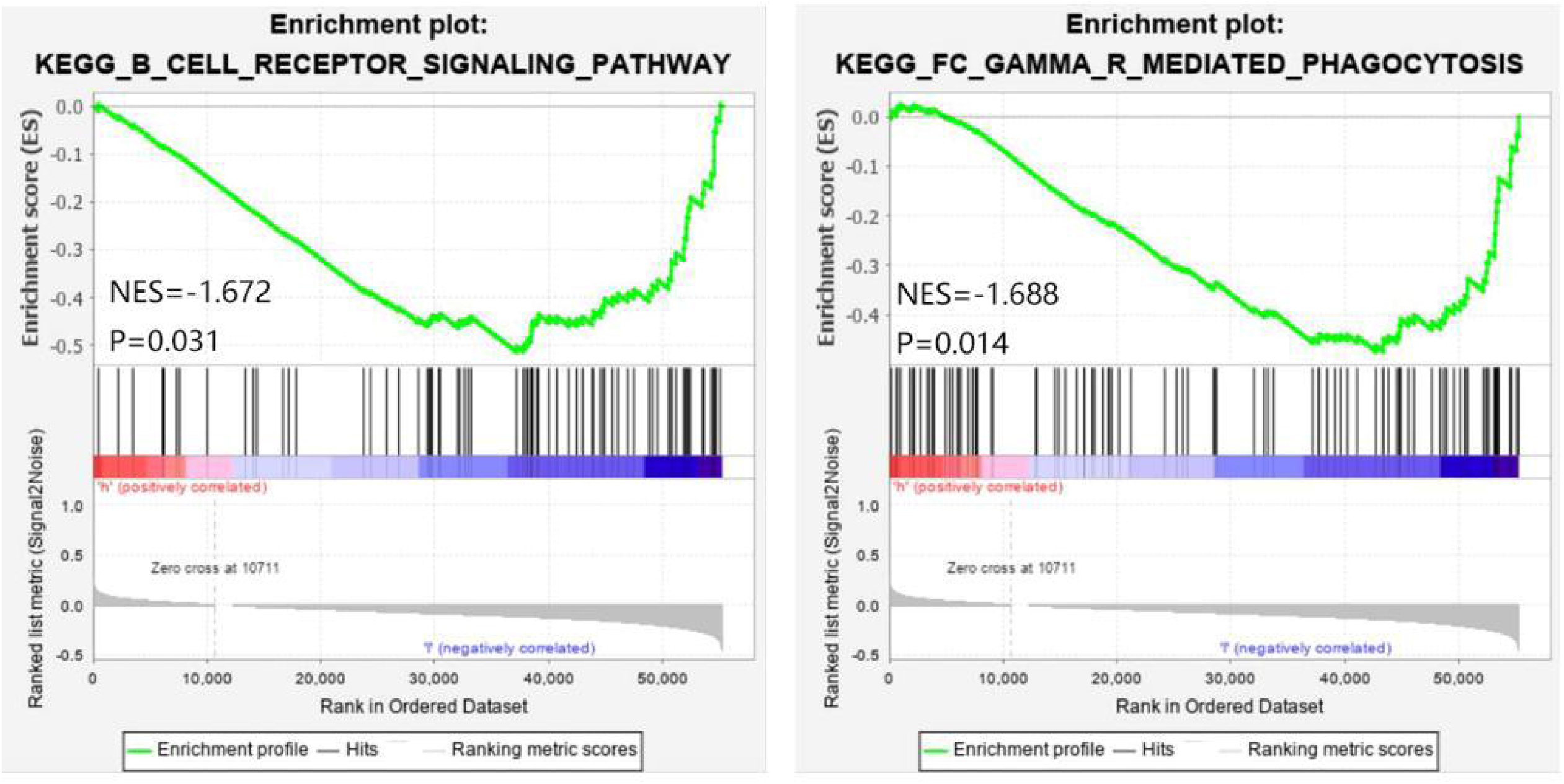
Results of gene set enrichment analysis (GSEA) in KEGG signaling pathway functional annotation

1. FC_GAMMA_R_MEDIATED_PHAGOCYTOSIS
2. B_CELL_RECEPTOR_SIGNALING_PATHWAY

#### Expression of model lncRNAs in different cell types

We analyzed gene distribution in the tumor single cell sequencing dataset GSE150321 in a 64-year-old male LSCC patient, which contained 3,885 single cell expression profiles. The results are shown in a scatter plot (Fig 10A). The distribution of lncRNA (AC104083.1, AC004148.2, AL357033.4, AC116914.2, AC024075.1, AC008115.3, PCED1B-AS1, RAB11B-AS1, AC004687.1, AL450992.2, EP300-AS1, AC024075.2 and AC136475.2) was further evaluated. Unfortunately, EP300-AS1 and AL450992.2 were not detected due to the restriction of sequencing depth. In accordance with the results of QRT-PCR, other lncRNA showed low expression in tumor. Interestingly, AC004687.1 and PCED1B-AS1 are highly expressed in T cells, but also in macrophages and dendritic cells (DC) (Fig 10B).

**Fig 10.**
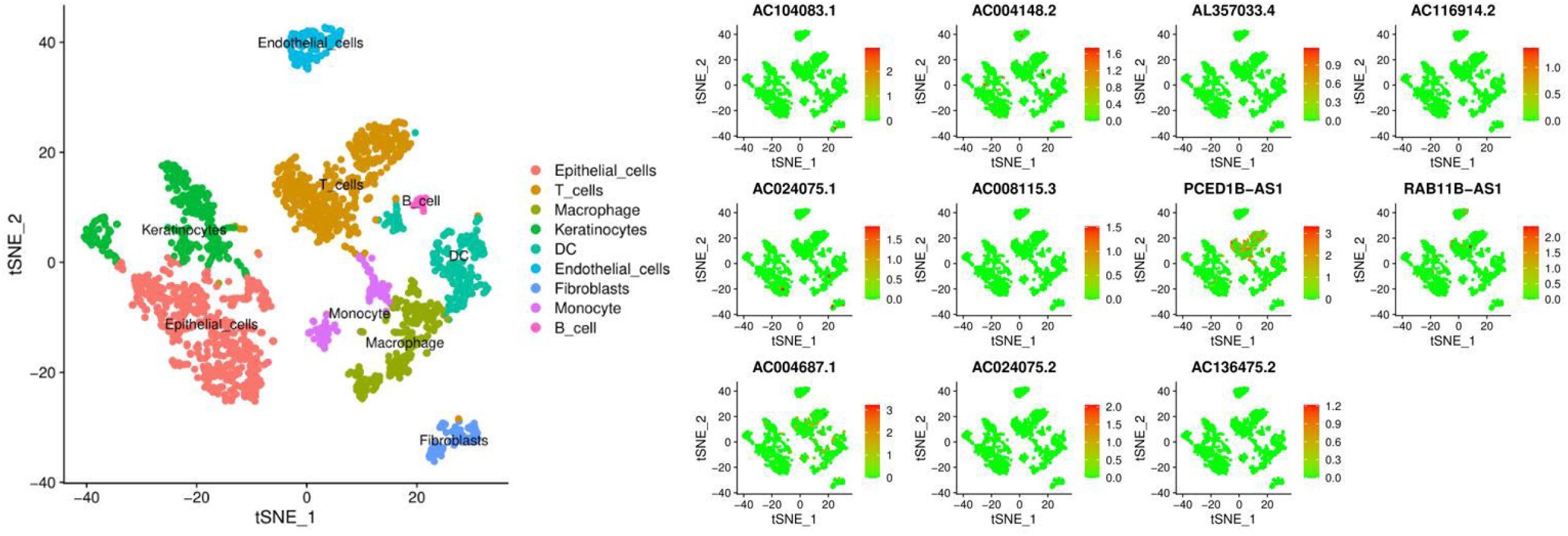
Single cell sequencing analysis. **(A)** Distribution of different cell types in LSCC. **(B)** Expression and distribution of model lncRNAs in 3,885 LSCC cells.

#### Function and signal pathway prediction of AC004687.1 and PCED1B-AS1

Furthermore, we screened out the protein coding genes associated with AC004687.1 and PCED1B-AS1. These genes were annotated with GO and KEGG analysis used to analyze the function and pathway of a single lncRNA. The results showed that AC004687.1 and PCED1B-AS1 related protein coding genes were involved in the immune process, (Fig 10, S3-6 Table, P<0.05). Seventeen enriched GO terms and 36 enriched KEGG pathways were associated with AC004687.1. The immune-related results of GO mainly including MHC protein complex binding, MHC class II protein complex binding and MHC protein binding, (Fig 11A, S3 Table). The immune-related results of significantly enriched pathways were Th17 cell differentiation, T cell receptor signaling pathway, natural killer cell mediated cytotoxicity, (Fig 11C, S5 Table). Eighty enriched GO terms and 54 enriched KEGG pathways were associated with PCED1B-AS1. Similarly, The results of GO mainly including MHC protein complex binding, immunoglobulin binding, MHC protein binding, MHC class II receptor activity, (Fig 11B, S4 Table). Significantly enriched pathways were Th1 and Th2 cell differentiation, T cell receptor signaling pathway, PD-L1 expression and PD-1 checkpoint pathway in cancer, (Fig 11D, S6 Table).

**Fig 11.**
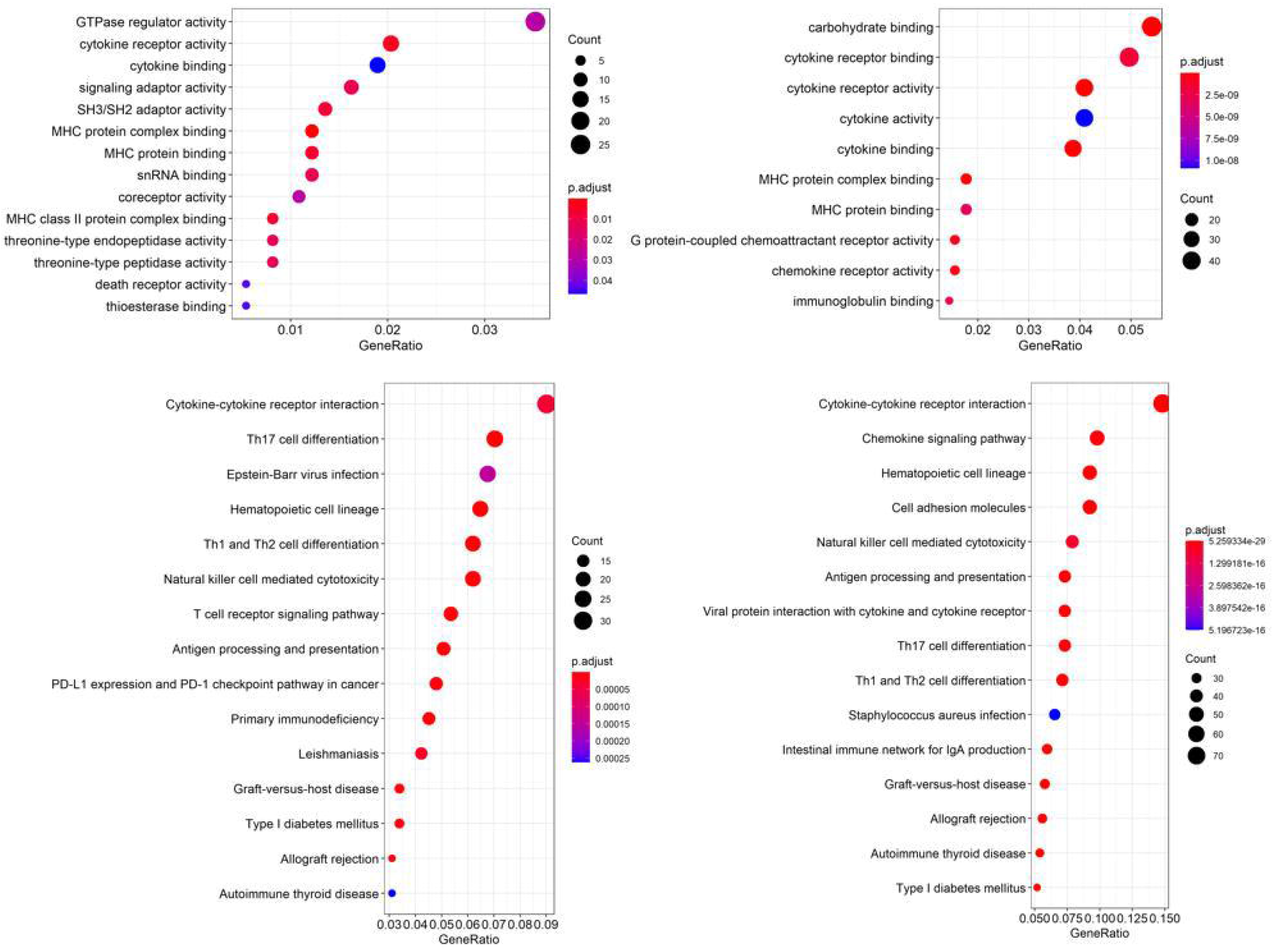
GO and KEGG pathway enrichment analysis for AC004687.1 and PCED1B-AS1 related mRNAs based on TCGA database. **(A)**Plot of the enriched GO terms associated with AC004687.1-related mRNAs. **(B)**Plot of the enriched GO terms associated with PCED1B-AS1-related mRNAs. **(C)**Plot of the enriched KEGG pathways associated with AC004687.1-related mRNAs. **(D)**Plot of the enriched KEGG pathways associated with PCED1B-AS1-related mRNAs.

Since both AC004687.1 and PCED1B-AS1 were significantly associated with Major histocompatibility complex type II (MHC II)receptor activity in GO functional annotation analysis, we detected the expression of MHC II classical molecules HLA-DRA and HLA-DRB1 in 17 HNSCC tissues and 9 normal tissues. And further analyzed the correlation between the expression of AC004687.1, PCED1B-AS1 and HLA-DRA, HLA-DRB1. The results showed that AC004687.1 was significant positively correlated with HLA-DRA (Pearson correlation coefficient=0.483, P=0.012). AC004687.1 was significant positively correlated with HLA-DRB1 (Pearson correlation coefficient=0.497, P=0.01). PCED1B-AS1 was significant positively correlated with HLA-DRA (Pearson correlation coefficient=0.550, P=0.004). PCED1B-AS1 was significant positively correlated with HLA-DRB1 (Pearson correlation coefficient=0.640,P=4.25E-04). (Fig 12A-D).

**Fig 12.**
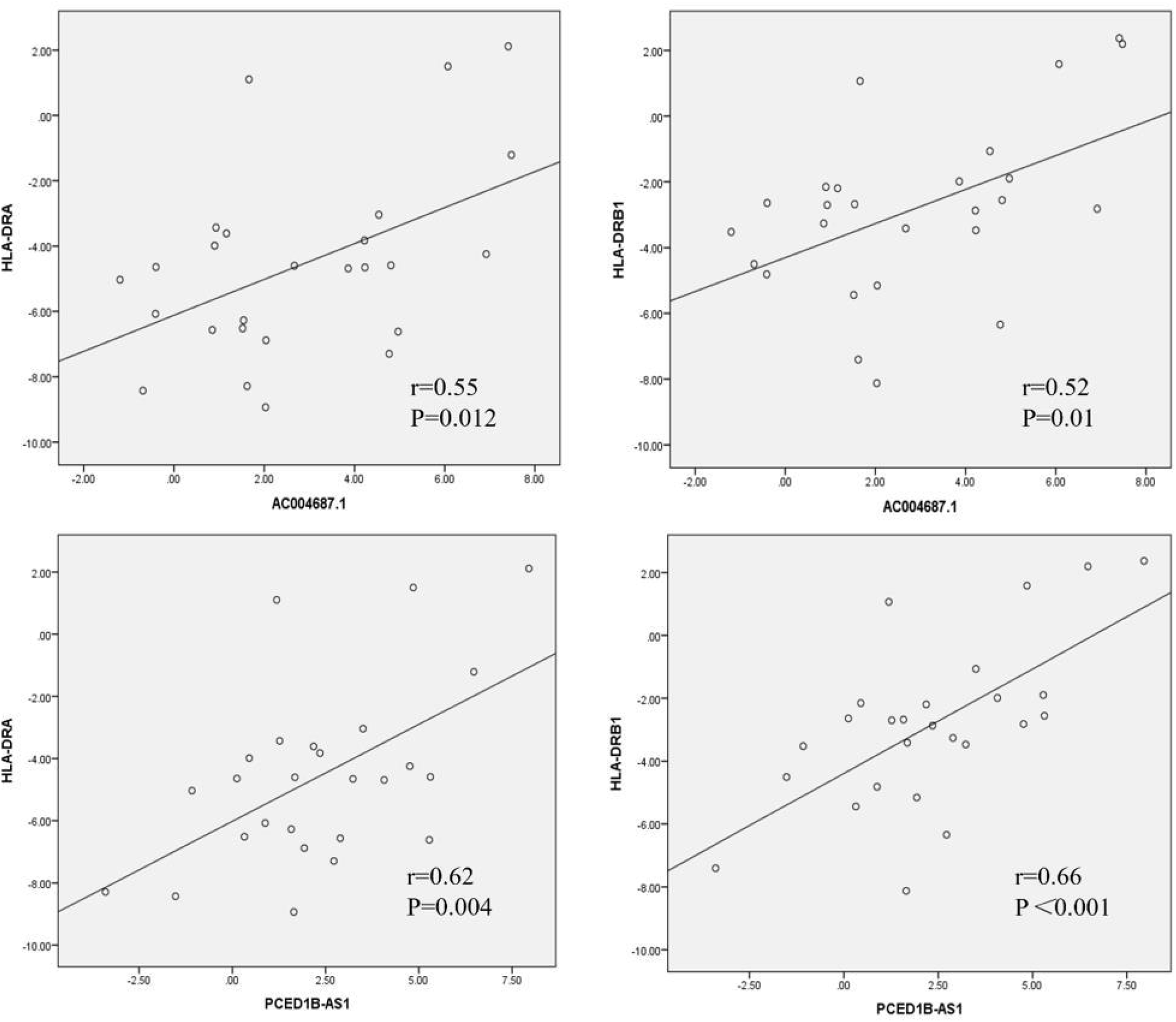
The correlation analysis of AC004687.1, PCED1B-AS1 and HLA-DRA, HLA-DRB1 expression in HNSCC tissue and normal tissue.

In order to further investigate the difference of HLA-DR expression at protein level in HNSCC tumor tissue samples, immunohistochemical staining and QRT-PCR were used to detect HLA-DR expression level and its protein distribution in 6 pairs of HNSCC tissues and their matched normal tissues. The results showed that HLA-DR protein staining mainly exists in the cell membrane and cytoplasm. The mean optical density (OD) in tumor samples were significantly lower than those in normal samples (P = 0.007). Similar to AC004687.1 and PCED1B-AS1, the expression of HLA-DR in tumor samples decreased significantly (Fig 13G, HLA-DRA: P=0.0315, HLA-DRB1: P=0.0207). The expression level in tumor matched normal tissue was significantly higher than that in tumor matched tissue. (Fig 13A-F)

**Fig 13.**
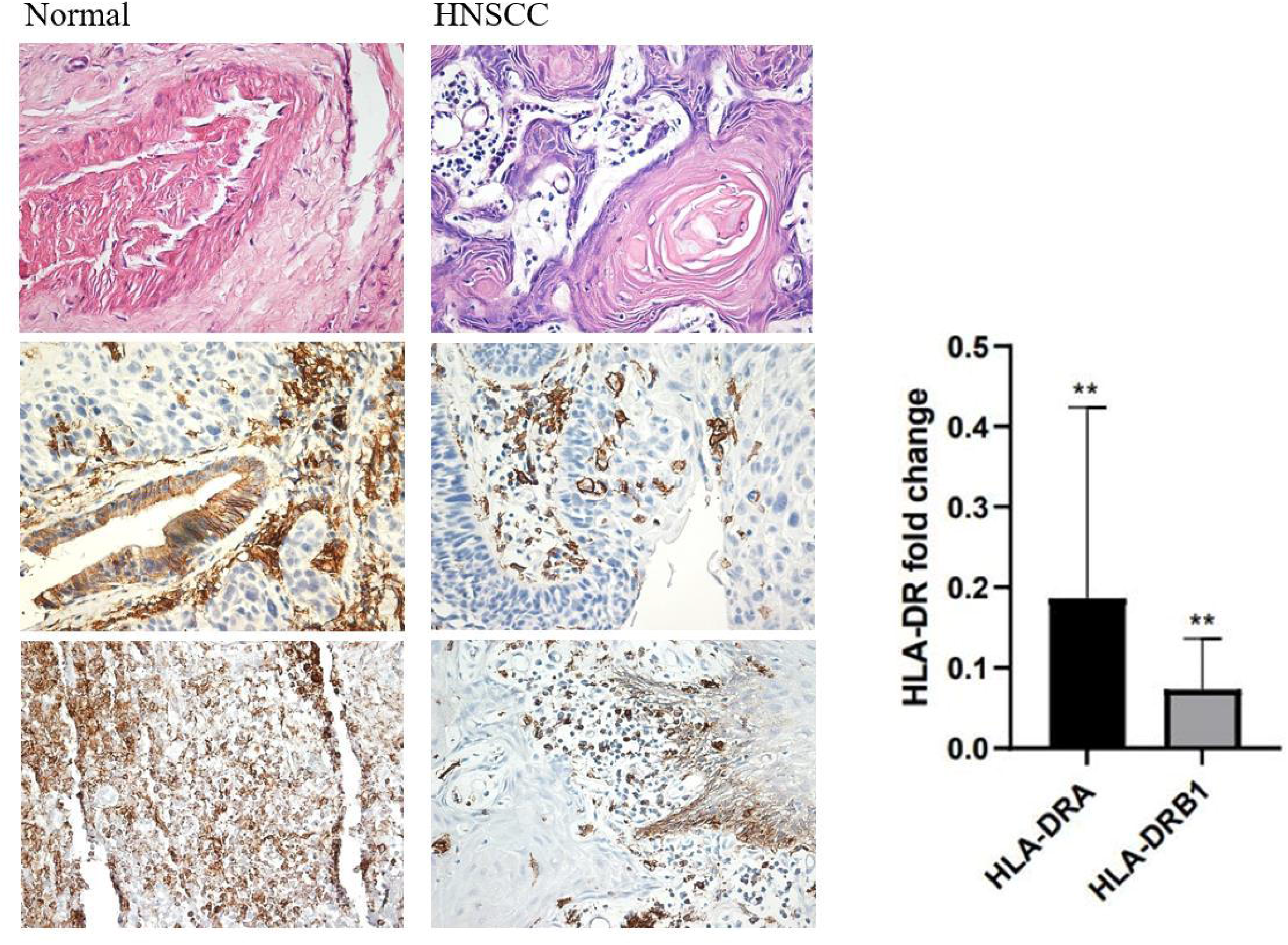
The expression of HLA-DR protein was down-regulated in HNSCC tumors. **(A, B)** HE staining of HNSCC tumor tissue and matched tissue adjacent to cancer. **(C-F)** HLA-DR immunohistochemical staining and DAB staining of HNSCC tumor tissue and matched tissue adjacent to cancer. **(G)** Differential expression of HLA-DR α and HLA-DR β1 in HNSCC tumors and matched tissues, *: P<0.05, **: P<0.01.

## DISCUSSION

In this research, we explored the immune-lncRNAs for the prognosis of head and neck squamous-cell carcinoma by transcriptional data and clinical information of 490 patients with HNSCC in The Cancer Genome Atlas (TCGA). According to coexpression and correlation analysis with immune gene sets from the Molecular Signatures Database (MSigDB) of GSEA (https://www.gsea-msigdb.org/gsea/msigdb/index.jsp), 17 immune-lncRNAs were identified to construct the prognostic risk model of HNSCC from 341 immune-related lncRNAs. The risk model was optimized based on whether individual lncRNA had survival prognostic significance and whether it was expressed differently in HNSCC and matched normal tissues. Single lncRNA survival analysis and QRT-PCR confirmed 13 immune-lncRNA including AL357033.4, AC024075.2, AC004687.1, AC024075.1, EP300-AS1, AL450992.2, AC008115.3, AC004148.2, RAB11B-AS1, AC136475.2, AC116914.2, PCED1B-AS1 and AC104083.1 showed difference in expression between tumor tissue and matched normal tissue, and the expression in normal tissue was higher than that in tumor tissue. Consistent with the results of previous Cox univariate analysis (Figure 1), which could be regarded as protective immunity-lncRNAs. Finally, 13 protective immune-lncRNAs used to construct the risk model. According to the median determined by the expression value of 13 immune-lncRNAs and the coefficient, patients were divided into a high-risk group and a low-risk group, and both groups underwent survival analysis. The results showed that the low-risk group had a five-year survival rate that was nearly two times greater than that of the high-risk group. The five-year survival rate of the high-risk group was approximately 35.2% (95% CI: 0.279-0.443), and that of the low-risk group was approximately 71.1% (95% CI: 0.630-0.803), P=1.923e-12. We further verified the independence and accuracy of the model, and the results of univariate and multivariate factor independent prognostic analysis showed that the risk score could be used as a high-risk factor to independently influence survival and prognosis (hazard ratio: 2.104、2.072, P<0.001). The risk score showed a higher AUC (0.701) than other clinical characteristics in the ROC curve. The results of the ROC curve and PCA showed that the high-risk group and the low-risk group had different immune statuses.

We further explored the results of KEGG signal pathway function annotation. Unfortunately, the high-risk group had no meaningful immune-related enrichment results. For the low-risk group, the main immune-related pathways were enriched in the Fc gamma R-mediated phagocytosis and B cell receptor signaling pathway. The phagocytosis mediated by Fc*γ*R showed that in the mechanism of antitumor activity mediated by monoclonal antibody, the Fc part of the antibody combined with activated Fc*γ* receptor (Fc*γ*R) showed greater inhibition of tumor growth and metastasis(35). As the main effector cells of humoral immunity, B cells can inhibit tumor progression by secreting immunoglobulin to promote T cell response and kill cancer cells(36, 37). The low-risk group may suppress the growth of the tumor by these cellular pathways.

By analyzing the laryngeal cancer single cell sequencing dataset GSE150321, we found that in immune-lncRNAs, AC004687.1 and PCED1B-AS1 were highly expressed in T cells, but also in macrophages and dendritic cells. There are few studies on PCED1B-AS1 in HNSCC. Consistent with our study, PCED1B-AS1 may play an important role in modulating the tumor immune microenvironment (TIME) and improving male HNSCC survival. We further explored their immune function. Interestingly, both AC004687.1 and PCED1B-AS1 showed significant associations with the MHC in the GO functional annotation. In response to MHC II receptor activity, both AC004687.1 and PCED1B-AS1 were significantly enriched, in this research, we detected the correlation between the expression of AC004687.1, PCED1B-AS1 and MHC II classical molecules HLA-DRA, HLA-DRB1. And we also detect the expression of HLA-DRA and HLA-DRB1 in HNSCC and adjacent tissues. They confirm that both AC004687.1 and PCED1B-AS1 are closely related to HLA-DR.

MHC II molecules stimulate CD4 + T cells and are active and key participants in anti-tumor immunity(38). HLA-DR as the MHC II antigen presenting molecule, expressed in the Antigen-presenting cell(39). It has been proved that the loss of HLA-DR expression is closely related to the occurrence and development of many kinds of tumors. HLA-DR has been reported to be an indicator of good prognosis in colorectal cancer, with survival times at least twice as long in patients with HLA-DR overexpression as in patients with low expression(40). Low expression of HLA-DR in epithelial cells of esophageal adenocarcinoma may be an independent predictor of low survival rate, with a 2.8-fold increase in disease-related mortality(41). For HNSCC, EGFR inhibitors have been reported to enhance HLA-DR expression in tumor cells as adjuvant therapy, thereby improving cancer immunotherapy(42). This research confirmed that lncRNAs AC004687.1 and PCED1B-AS1 were positively correlated with HLA-DRA and HLA-DRB1 expression in HNSCC, the effect of HLA-DR on head and neck squamous-cell carcinoma remains to be studied. AC004687.1 and PCED1B-AS1 as protective lncRNA were significantly correlated with T stage in the correlation analysis of clinical characteristics, (AC004687.1:P<0.001,PCED1B-AS1:P <0.05), the expression decreased with the increase of T. Therefore, we reasonably infer that AC004687.1 and PCED1B-AS1 through HLA-DR participating in MHC class II immune pathway exert anti-tumor effect.

Multiple RNAs (encoding and noncoding RNAs) of HNSCC patients have abnormal changes and play key roles in cancer biology, and long noncoding RNAs can be used as biomarkers for predicting therapy or diagnostic tools(43). However, the data of specific lncRNAs for HNSCC are scarce(44), and there is little research on immune-related lncRNAs. We hope to provide a reference for HNSCC targeted therapy from the point of immune correlation by data analysis of bioinformatics. Compared with the previous immune-associated lncRNA risk model(21), the conditions for constructing the model were optimized by experimental analysis and the HR of the risk model in this study was more than 2(45). In addition, we have further explored the distribution of these lncRNAs in LSCC cells and the immune function. However, we need further research to explore the regulatory mechanism and detailed relationship between the expression of lncRNA and MHC II. The function of unannotated lncRNAs for tumor occurrence and development, such as AC004687.1, showed a significant correlation with T in the correlation analysis of clinical characteristics(P=3.6e−05), therefore, AC004687.1 merits further research.

## CONCLUSION

Our study established a risk model constructed from 13 immune-lncRNA with differential expression, which can be used to differentiate the immune status of HNSSC patients and predict their survival prognosis. AC004687.1 and PCED1B-AS1 were overexpressed on T cells in LSCC, participated in immune response through MHC II pathway, which can be used as a marker of good prognosis in HNSCC patients and as a potential therapeutic target provide reference for specific immunotherapy in HNSCC patients.

## Acknowledgments

The authors thank all the participants in this study for their support.

## Supporting information

S1 Table. The clinicopathological information of 17 HNSCC patients

S2 Table. List of primers used for QRT-PCR

S1 Fig. Single-factor significant immune-lncRNAs.

A hazard ratio >1 is a high-risk factor, and a hazard ratio <1 is a low-risk factor.

S2 Fig. Survival analysis of single immune-lncRNAs.

S3 Fig. GSEA indicated more active trends in the enrichment of the immune phenotype in the low-risk group.

S3 Table. The GO enrichment analysis for AC004687.1-related mRNAs

S4 Table. The GO enrichment analysis for PCED1B-AS1-related mRNAs.

S5 Table. The KEGG pathway enrichment analysis for AC004687.1-related mRNAs

S6 Table. The KEGG pathway enrichment analysis for PCED1B-AS1-related mRNA.

---

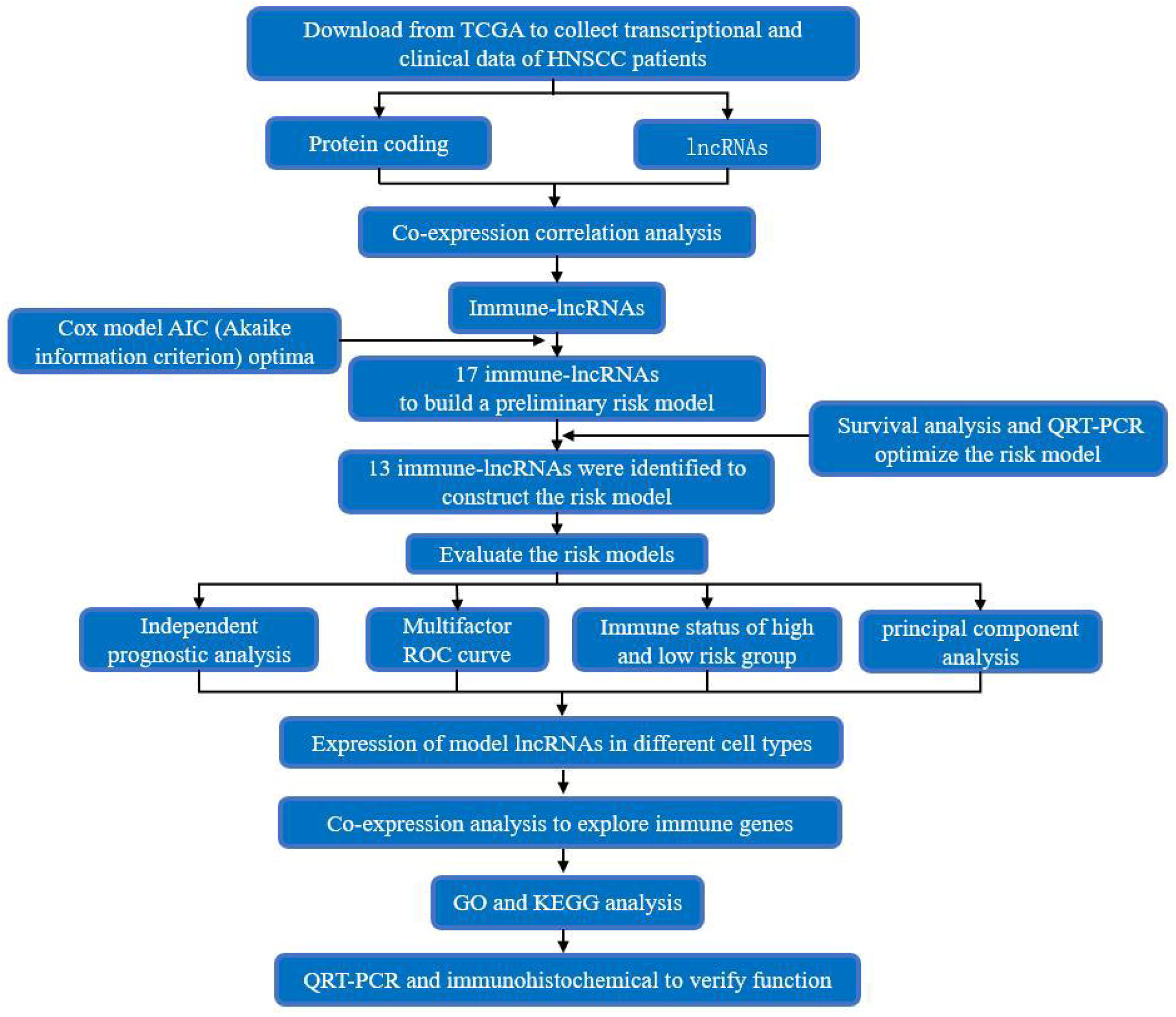

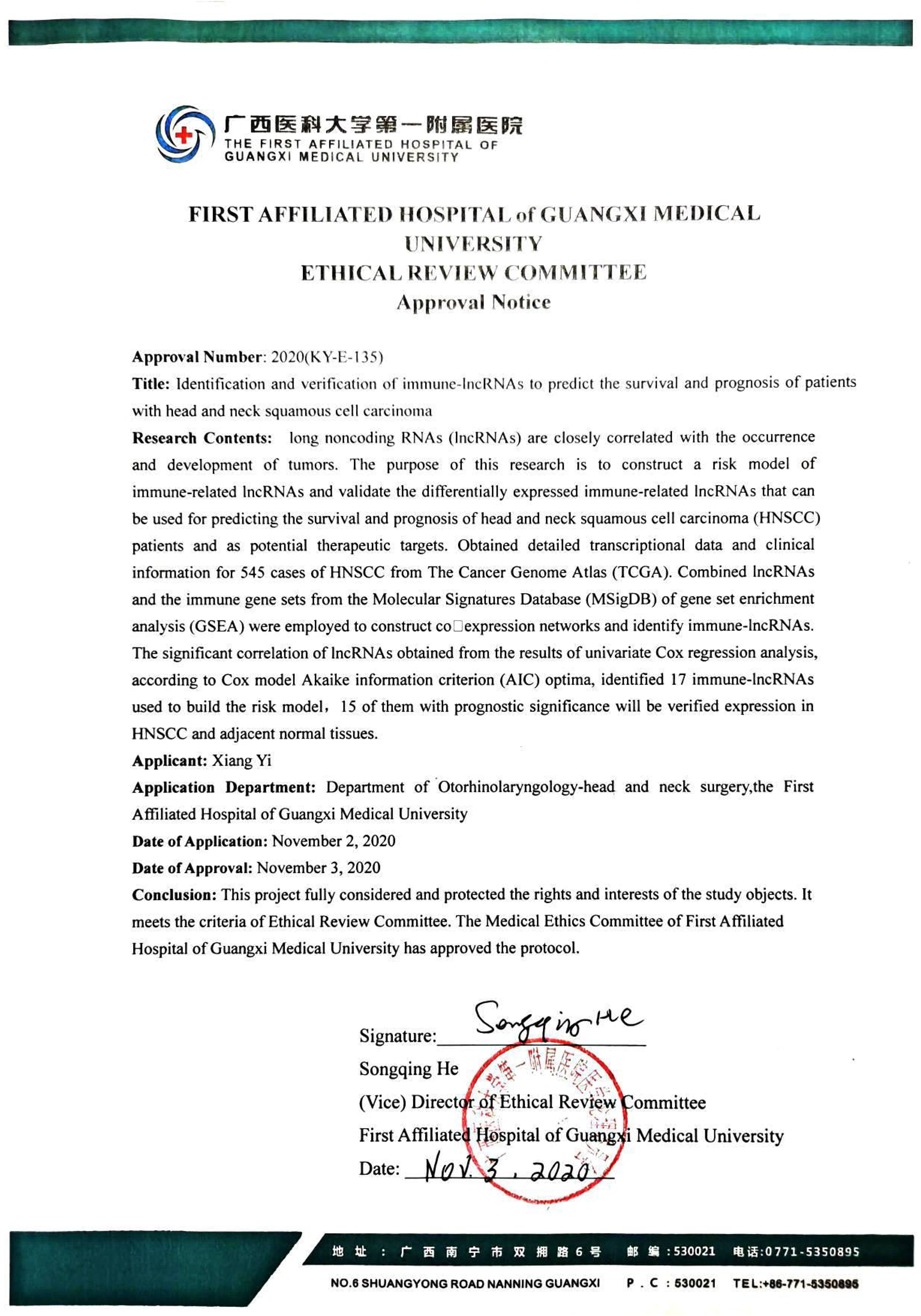

