## Supplemental 1 for "An immune-lncRNA risk model to predict prognosis for patients with head and neck squamous cell carcinoma": Supplemental 1.docx

| No. | Sex | Age (years) | Diagnosis | Clinical stage | Histological grade |
| --- | --- | --- | --- | --- | --- |
| 1 | M | 58 | HSCC | T2N0M0 | Moderate |
| 2 | M | 31 | HSCC | T4aN2M0 | Well |
| 3 | M | 55 | LSCC | T4N0M0 | Well |
| 4 | M | 66 | LSCC | T4N2M0 | Well |
| 5 | M | 45 | HSCC | T2N2M0 | Well |
| 6 | M | 64 | HSCC | T4N0M0 | Well/Moderate |
| 7 | M | 52 | TSCC | T4N2M0 | Well/Moderate |
| 8 | M | 69 | TSCC | T2N2M0 | Moderate |
| 9 | M | 70 | LSCC | T1aN0M0 | Well/Moderate |
| 10 | F | 54 | TSCC | T4N0M0 | Well/Moderate |
| 11 | M | 69 | LSCC | T3N0M0 | Well |
| 12 | M | 56 | HSCC | T4aN2M0 | Well/Moderate |
| 13 | M | 56 | HSCC | T2N1M0 | Moderate |
| 14 | M | 57 | LSCC | T4N0Mx | Well |
| 15 | M | 54 | TSCC | T1N2M0 | Well |
| 16 | M | 59 | LSCC | T4N1M0 | Well/Moderate |
| 17 | M | 73 | LSCC | T2NxM0 | Moderate |

**S1 Table.** The clinicopathological information of 17 HNSCC patients

M, male; F, female; TSCC, tongue squamous cell carcinoma; HSCC, hypopharyngeal squamous-cell carcinoma, LSCC, Laryngeal squamous-cell carcinoma.

| Genes | Sequences (5’-3’) |
| --- | --- |
| HLA-DRA | Forward primer: 5’- GCCGAGTTCTATCTGAATCCTGACC-3’ |
|  | Reverse primer: 5’- GACCGTCTCCTTCTTTGCCATATCC-3’ |
| HLA-DRB1 | Forward primer: 5’- GGAATGGAGAGCACGGTCTGAATC-3’ |
|  | Reverse primer: 5’- GGCTGAAGTCCAGAGTGTCCTTTC-3’ |
| GAPDH | Forward primer: 5’-ACAACTTTGGTATCGTGGAAGG-3’ |
|  | Reverse primer: 5’-GCCATCACGCCACAGTTTC-3’ |
| IER3-AS1 | Forward primer:5’-TCTACCTCGCAGCCACCCTAAAG-3’ |
|  | Reverse primer: 5’-GCATCCTCCAGCATCTCAACTCC-3’ |
| AL357033.4 | Forward primer: 5’-TCAGCATCCCAGGTCGAGTC-3’ |
|  | Reverse primer: 5’-ACCATCGCACATAGCGTCAG-3’ |
| EP300-AS1 | Forward primer:5’-TCAAGCCTTCATTTCCCTCTCAA-3’ |
|  | Reverse primer: 5’-TCCTCTCAGGGTTTGTCCAATG-3’ |
| PCED1B-AS1 | Forward primer:5’-GCAGGCTGAGGAATTACCAAGGAG-3’ |
|  | Reverse primer: 5’-GGAGGGCACAAGATCAGATGAACC-3’ |
| AC004687.1 | Forward primer: 5’-CAGCCTGAAGAGTACACGCC-3’ |
|  | Reverse primer: 5’-TCTCCGAAGCCCACAGTACA-3’ |
| AC024075.2 | Forward primer: 5’-ACCCGCCTACCTCAGCCTTTC-3’ |
|  | Reverse primer: 5’-AGTTACCACACAGCCCAGCAATTC-3’ |
| AC024075.1 | Forward primer: 5’-TCAGAACAGCAGAACAGCAGC -3’ |
|  | Reverse primer: 5’-TTGCTAATGACAGCTTCGCTGG-3’ |
| AC015849.3 | Forward primer:5’-CTCCCACCTCAGCCTCCTAAGTAG-3’ |
|  | Reverse primer: 5’-AGTGAGCCGAGATTGCACCATTG-3’ |
| AC116914.2 | Forward primer:5’-GCCTGTAATCCCAGCACTTCGG-3’ |
|  | Reverse primer: 5’-GCAGTGGCACGATCTCTACTCAC-3’ |
| RAB11B-AS1 | Forward primer: 5’-ATGTTGGCCCGGCTGGTCTC-3’ |
|  | Reverse primer: 5’-CGGTGTCTCGTGCCTGTAATCC-3’ |
| AC104083.1 | Forward primer:5’-GCTACAGTTGGCTAGTGGGTCTTC-3’ |
|  | Reverse primer: 5’-AGCTCTGGCAGTACATGTGTCTTG-3’ |
| AC004148.2 | Forward primer: 5’-CCACTTGAGCACCGCCACATC-3’ |
|  | Reverse primer: 5’-CCAGTGAGGTCAGGACGTAGGG-3’ |
| AC008115.3 | Forward primer:5’-AATGGGGAAGGCTGGGGATACG-3’ |
|  | Reverse primer: 5’-TAGTGGGACTCACAGGGAAACGAC-3’ |
| AL450992.2 | Forward primer:5’-ACCCACTGCACCCTCAACTC-3’ |
|  | Reverse primer: 5’-TCCCCTGAACCACCTCCTCA-3’ |
| AC136475.2 | Forward primer:5’-AGCACTGACGTTCCCGGAAA-3’ |
|  | Reverse primer: 5’-TTCTGGTTTCGGGCCTCCTG-3’ |

**S2 Table.** List of primers used for QRT-PCR

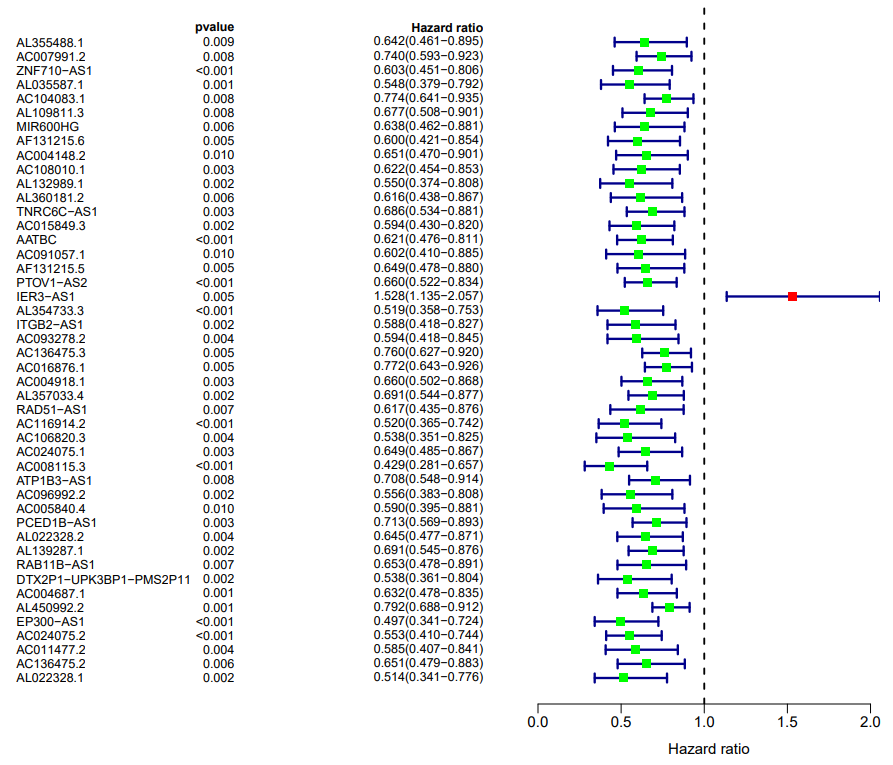

**S1 Fig.** Single-factor significant immune-lncRNAs.

A hazard ratio >1 is a high-risk factor, and a hazard ratio <1 is a low-risk factor.

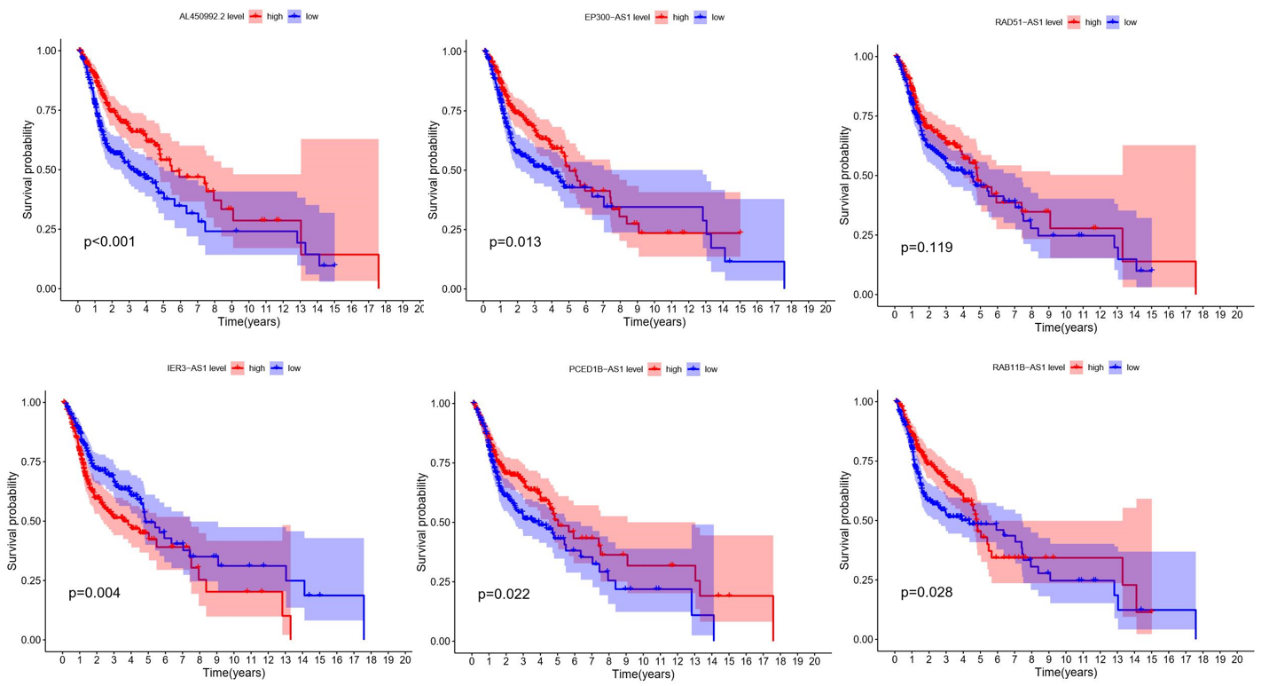

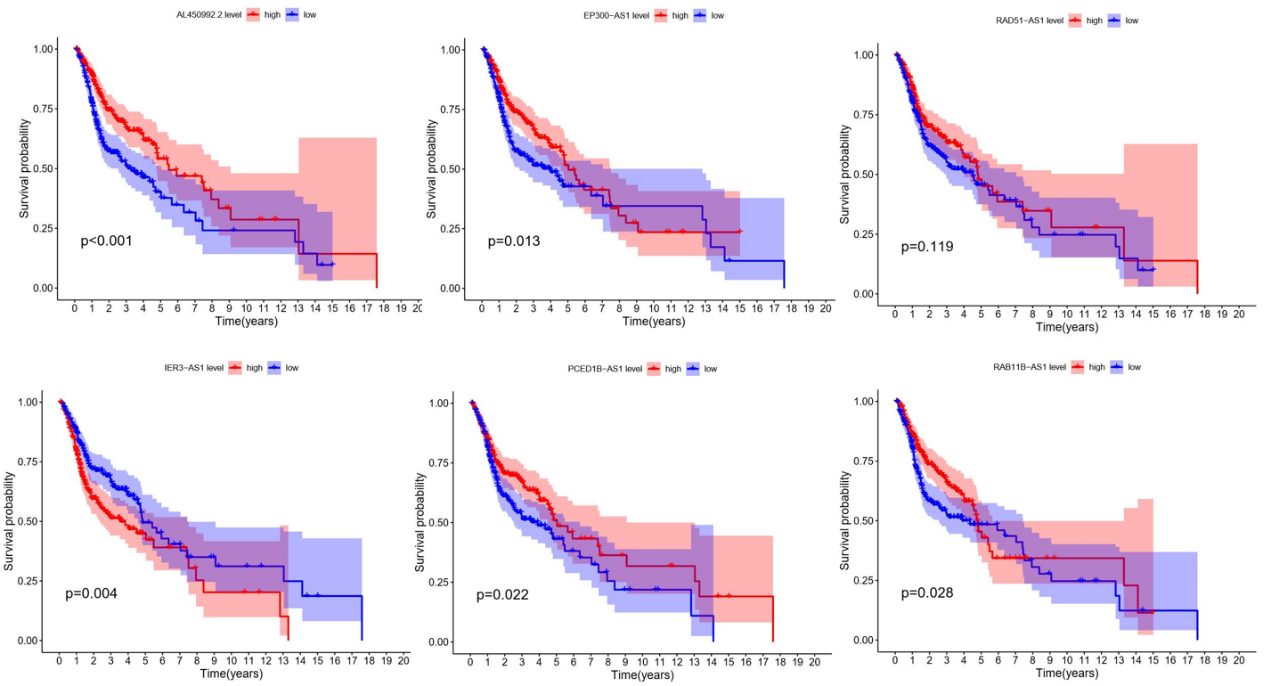

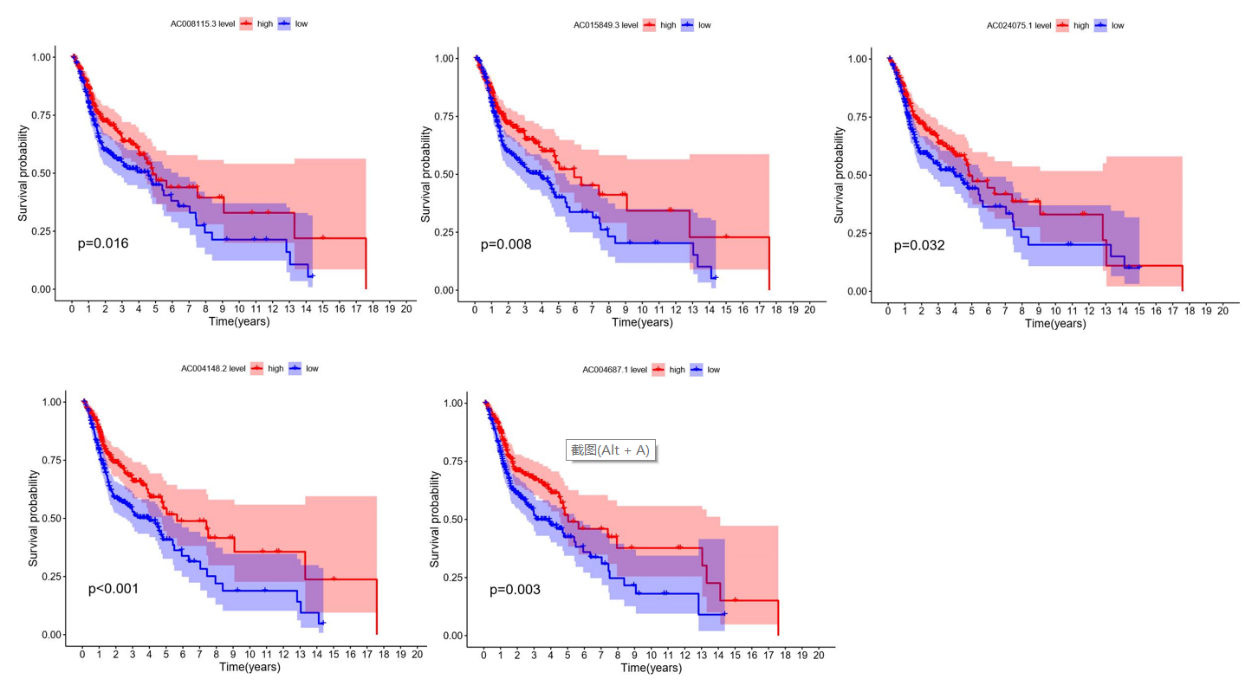
 **S2 Fig.** Survival analysis of single immune-lncRNAs.

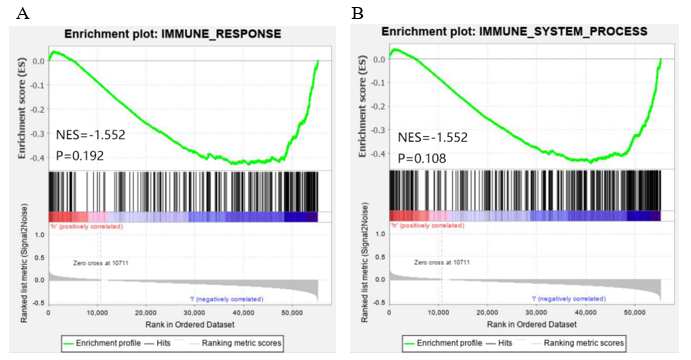
 **S3 Fig.** GSEA indicated more active trends in the enrichment of the immune phenotype in the low-risk group.

| ID | Description | GeneRatio | BgRatio | Pvalue | P.adjust | Qvalue | Count |
| --- | --- | --- | --- | --- | --- | --- | --- |
| GO:0023023 | MHC protein complex binding | 9/738 | 25/17697 | 4.07E-07 | 0.000292 | 0.000277 | 9 |
| GO:0004896 | cytokine receptor activity | 15/738 | 96/17697 | 9.80E-06 | 0.003514 | 0.003338 | 15 |
| GO:0023026 | MHC class II protein complex binding | 6/738 | 16/17697 | 2.88E-05 | 0.005563 | 0.005284 | 6 |
| GO:0042287 | MHC protein binding | 9/738 | 40/17697 | 3.10E-05 | 0.005563 | 0.005284 | 9 |
| GO:0005070 | SH3/SH2 adaptor activity | 10/738 | 52/17697 | 4.83E-05 | 0.006927 | 0.00658 | 10 |
| GO:0035591 | signaling adaptor activity | 12/738 | 80/17697 | 0.000113 | 0.0122 | 0.011588 | 12 |
| GO:0017069 | snRNA binding | 9/738 | 47/17697 | 0.000119 | 0.0122 | 0.011588 | 9 |
| GO:0004298 | threonine-type endopeptidase activity | 6/738 | 21/17697 | 0.000163 | 0.013009 | 0.012357 | 6 |
| GO:0070003 | threonine-type peptidase activity | 6/738 | 21/17697 | 0.000163 | 0.013009 | 0.012357 | 6 |
| GO:0015026 | coreceptor activity | 8/738 | 44/17697 | 0.000412 | 0.028873 | 0.027425 | 8 |
| GO:0030695 | GTPase regulator activity | 26/738 | 304/17697 | 0.000443 | 0.028873 | 0.027425 | 26 |
| GO:0005035 | death receptor activity | 4/738 | 11/17697 | 0.000783 | 0.043168 | 0.041004 | 4 |
| GO:0031996 | thioesterase binding | 4/738 | 11/17697 | 0.000783 | 0.043168 | 0.041004 | 4 |
| GO:0019955 | cytokine binding | 14/738 | 128/17697 | 0.000911 | 0.046672 | 0.044332 | 14 |

**S3 Table.** The GO enrichment analysis for AC004687.1-related mRNAs

| ID | Description | GeneRatio | BgRatio | Pvalue | P.adjust | Qvalue | Count |
| --- | --- | --- | --- | --- | --- | --- | --- |
| GO:0004896 | cytokine receptor activity | 37/905 | 96/17697 | 2.37E-23 | 1.83E-20 | 1.61E-20 | 37 |
| GO:0019955 | cytokine binding | 35/905 | 128/17697 | 1.14E-16 | 4.40E-14 | 3.88E-14 | 35 |
| GO:0023023 | MHC protein complex binding | 16/905 | 25/17697 | 2.55E-15 | 6.55E-13 | 5.77E-13 | 16 |
| GO:0030246 | carbohydrate binding | 49/905 | 271/17697 | 7.82E-15 | 1.51E-12 | 1.33E-12 | 49 |
| GO:0001637 | G protein-coupled chemoattractant receptor activity | 14/905 | 26/17697 | 4.12E-12 | 5.30E-10 | 4.67E-10 | 14 |
| GO:0004950 | chemokine receptor activity | 14/905 | 26/17697 | 4.12E-12 | 5.30E-10 | 4.67E-10 | 14 |
| GO:0005126 | cytokine receptor binding | 45/905 | 286/17697 | 1.51E-11 | 1.66E-09 | 1.46E-09 | 45 |
| GO:0019865 | immunoglobulin binding | 13/905 | 24/17697 | 2.22E-11 | 2.14E-09 | 1.89E-09 | 13 |
| GO:0042287 | MHC protein binding | 16/905 | 40/17697 | 3.79E-11 | 3.25E-09 | 2.86E-09 | 16 |
| GO:0005125 | cytokine activity | 37/905 | 220/17697 | 1.41E-10 | 1.09E-08 | 9.61E-09 | 37 |
| GO:0015026 | coreceptor activity | 16/905 | 44/17697 | 2.07E-10 | 1.45E-08 | 1.28E-08 | 16 |
| GO:0016493 | C-C chemokine receptor activity | 12/905 | 23/17697 | 2.39E-10 | 1.54E-08 | 1.35E-08 | 12 |
| GO:0019957 | C-C chemokine binding | 12/905 | 24/17697 | 4.55E-10 | 2.70E-08 | 2.38E-08 | 12 |
| GO:0035586 | purinergic receptor activity | 12/905 | 25/17697 | 8.35E-10 | 4.61E-08 | 4.06E-08 | 12 |
| GO:0042605 | peptide antigen binding | 13/905 | 31/17697 | 1.31E-09 | 6.76E-08 | 5.95E-08 | 13 |
| GO:0032395 | MHC class II receptor activity | 8/905 | 10/17697 | 1.86E-09 | 8.99E-08 | 7.92E-08 | 8 |
| GO:0003823 | antigen binding | 29/905 | 160/17697 | 2.36E-09 | 1.07E-07 | 9.45E-08 | 29 |
| GO:0004715 | non-membrane spanning protein tyrosine kinase activity | 15/905 | 46/17697 | 4.39E-09 | 1.88E-07 | 1.66E-07 | 15 |
| GO:0001614 | purinergic nucleotide receptor activity | 10/905 | 20/17697 | 1.35E-08 | 4.95E-07 | 4.36E-07 | 10 |
| GO:0016502 | nucleotide receptor activity | 10/905 | 20/17697 | 1.35E-08 | 4.95E-07 | 4.36E-07 | 10 |
| GO:0042288 | MHC class I protein binding | 10/905 | 20/17697 | 1.35E-08 | 4.95E-07 | 4.36E-07 | 10 |
| GO:0019956 | chemokine binding | 12/905 | 32/17697 | 2.60E-08 | 9.12E-07 | 8.04E-07 | 12 |
| GO:0005070 | SH3/SH2 adaptor activity | 15/905 | 52/17697 | 2.88E-08 | 9.68E-07 | 8.53E-07 | 15 |
| GO:0001664 | G protein-coupled receptor binding | 38/905 | 280/17697 | 3.78E-08 | 1.22E-06 | 1.07E-06 | 38 |
| GO:0035591 | signaling adaptor activity | 18/905 | 80/17697 | 8.52E-08 | 2.63E-06 | 2.32E-06 | 18 |
| GO:0008009 | chemokine activity | 14/905 | 49/17697 | 9.56E-08 | 2.84E-06 | 2.50E-06 | 14 |
| GO:0048020 | CCR chemokine receptor binding | 13/905 | 43/17697 | 1.31E-07 | 3.75E-06 | 3.31E-06 | 13 |
| GO:0042379 | chemokine receptor binding | 16/905 | 66/17697 | 1.47E-07 | 4.06E-06 | 3.58E-06 | 16 |
| GO:0005164 | tumor necrosis factor receptor binding | 11/905 | 31/17697 | 1.94E-07 | 5.17E-06 | 4.55E-06 | 11 |
| GO:0019864 | IgG binding | 7/905 | 11/17697 | 2.46E-07 | 6.34E-06 | 5.58E-06 | 7 |
| GO:0008528 | G protein-coupled peptide receptor activity | 24/905 | 146/17697 | 3.75E-07 | 9.34E-06 | 8.23E-06 | 24 |
| GO:0023026 | MHC class II protein complex binding | 8/905 | 16/17697 | 4.04E-07 | 9.76E-06 | 8.59E-06 | 8 |
| GO:0001653 | peptide receptor activity | 24/905 | 152/17697 | 8.00E-07 | 1.87E-05 | 1.65E-05 | 24 |
| GO:0001608 | G protein-coupled nucleotide receptor activity | 7/905 | 13/17697 | 1.17E-06 | 2.58E-05 | 2.27E-05 | 7 |
| GO:0045028 | G protein-coupled purinergic nucleotide receptor activity | 7/905 | 13/17697 | 1.17E-06 | 2.58E-05 | 2.27E-05 | 7 |
| GO:0071723 | lipopeptide binding | 6/905 | 10/17697 | 3.09E-06 | 6.64E-05 | 5.85E-05 | 6 |
| GO:0042277 | peptide binding | 34/905 | 295/17697 | 8.30E-06 | 0.000173 | 0.000152 | 34 |
| GO:0004126 | cytidine deaminase activity | 6/905 | 12/17697 | 1.25E-05 | 0.000253 | 0.000223 | 6 |
| GO:0032813 | tumor necrosis factor receptor superfamily binding | 11/905 | 46/17697 | 1.51E-05 | 0.000299 | 0.000263 | 11 |
| GO:0001618 | virus receptor activity | 14/905 | 74/17697 | 1.96E-05 | 0.000369 | 0.000325 | 14 |
| GO:0104005 | hijacked molecular function | 14/905 | 74/17697 | 1.96E-05 | 0.000369 | 0.000325 | 14 |
| GO:0038024 | cargo receptor activity | 15/905 | 85/17697 | 2.38E-05 | 0.000438 | 0.000386 | 15 |
| GO:0033218 | amide binding | 37/905 | 356/17697 | 3.45E-05 | 0.00062 | 0.000546 | 37 |
| GO:0048018 | receptor ligand activity | 46/905 | 482/17697 | 3.57E-05 | 0.000626 | 0.000551 | 46 |
| GO:0001848 | complement binding | 7/905 | 21/17697 | 5.52E-05 | 0.000927 | 0.000817 | 7 |
| GO:0038187 | pattern recognition receptor activity | 7/905 | 21/17697 | 5.52E-05 | 0.000927 | 0.000817 | 7 |
| GO:0042169 | SH2 domain binding | 9/905 | 36/17697 | 6.20E-05 | 0.001018 | 0.000897 | 9 |
| GO:0030695 | GTPase regulator activity | 32/905 | 304/17697 | 9.01E-05 | 0.001449 | 0.001276 | 32 |
| GO:0019239 | deaminase activity | 8/905 | 32/17697 | 0.000159 | 0.002507 | 0.002208 | 8 |
| GO:0033691 | sialic acid binding | 5/905 | 12/17697 | 0.000203 | 0.003129 | 0.002756 | 5 |
| GO:0016814 | hydrolase activity, acting on carbon-nitrogen (but not peptide) bonds, in cyclic amidines | 8/905 | 34/17697 | 0.000251 | 0.003796 | 0.003344 | 8 |
| GO:0005085 | guanyl-nucleotide exchange factor activity | 24/905 | 214/17697 | 0.000258 | 0.003836 | 0.003378 | 24 |
| GO:0005068 | transmembrane receptor protein tyrosine kinase adaptor activity | 5/905 | 13/17697 | 0.000315 | 0.004595 | 0.004047 | 5 |
| GO:0008329 | signaling pattern recognition receptor activity | 6/905 | 20/17697 | 0.000367 | 0.00521 | 0.004589 | 6 |
| GO:0005096 | GTPase activator activity | 28/905 | 273/17697 | 0.000371 | 0.00521 | 0.004589 | 28 |
| GO:0030674 | protein binding, bridging | 20/905 | 170/17697 | 0.000442 | 0.006088 | 0.005363 | 20 |
| GO:0046935 | 1-phosphatidylinositol-3-kinase regulator activity | 5/905 | 14/17697 | 0.00047 | 0.006367 | 0.005608 | 5 |
| GO:0004713 | protein tyrosine kinase activity | 17/905 | 134/17697 | 0.000486 | 0.006469 | 0.005698 | 17 |
| GO:0001540 | amyloid-beta binding | 12/905 | 78/17697 | 0.000572 | 0.007483 | 0.006591 | 12 |
| GO:0043548 | phosphatidylinositol 3-kinase binding | 7/905 | 30/17697 | 0.000646 | 0.008313 | 0.007322 | 7 |
| GO:0001846 | opsonin binding | 5/905 | 15/17697 | 0.000676 | 0.00855 | 0.007531 | 5 |
| GO:0048365 | Rac GTPase binding | 11/905 | 69/17697 | 0.000706 | 0.008788 | 0.007741 | 11 |
| GO:0001784 | phosphotyrosine residue binding | 8/905 | 40/17697 | 0.000809 | 0.009891 | 0.008712 | 8 |
| GO:0060589 | nucleoside-triphosphatase regulator activity | 32/905 | 344/17697 | 0.00082 | 0.009891 | 0.008712 | 32 |
| GO:0046625 | sphingolipid binding | 6/905 | 23/17697 | 0.000838 | 0.009958 | 0.008771 | 6 |
| GO:1990782 | protein tyrosine kinase binding | 13/905 | 93/17697 | 0.000875 | 0.010237 | 0.009017 | 13 |
| GO:0017124 | SH3 domain binding | 16/905 | 130/17697 | 0.000981 | 0.011304 | 0.009957 | 16 |
| GO:0045309 | protein phosphorylated amino acid binding | 9/905 | 51/17697 | 0.001007 | 0.011431 | 0.010069 | 9 |
| GO:0035014 | phosphatidylinositol 3-kinase regulator activity | 5/905 | 17/17697 | 0.001278 | 0.014096 | 0.012416 | 5 |
| GO:0050664 | oxidoreductase activity, acting on NAD(P)H, oxygen as acceptor | 5/905 | 17/17697 | 0.001278 | 0.014096 | 0.012416 | 5 |
| GO:0031996 | thioesterase binding | 4/905 | 11/17697 | 0.001679 | 0.018254 | 0.016079 | 4 |
| GO:0019903 | protein phosphatase binding | 16/905 | 140/17697 | 0.002148 | 0.023027 | 0.020283 | 16 |
| GO:0016004 | phospholipase activator activity | 4/905 | 12/17697 | 0.002417 | 0.02556 | 0.022514 | 4 |
| GO:0060090 | molecular adaptor activity | 23/905 | 237/17697 | 0.002462 | 0.025684 | 0.022623 | 23 |
| GO:0008081 | phosphoric diester hydrolase activity | 12/905 | 94/17697 | 0.002984 | 0.030674 | 0.027019 | 12 |
| GO:0019902 | phosphatase binding | 19/905 | 185/17697 | 0.00302 | 0.030674 | 0.027019 | 19 |
| GO:0005044 | scavenger receptor activity | 8/905 | 51/17697 | 0.004084 | 0.040947 | 0.036067 | 8 |
| GO:0017048 | Rho GTPase binding | 18/905 | 177/17697 | 0.004253 | 0.042095 | 0.037078 | 18 |
| GO:0060229 | lipase activator activity | 4/905 | 14/17697 | 0.004503 | 0.044007 | 0.038762 | 4 |
| GO:0043028 | cysteine-type endopeptidase regulator activity involved in apoptotic process | 7/905 | 42/17697 | 0.005025 | 0.048494 | 0.042715 | 7 |

**S4 Table.** The GO enrichment analysis for PCED1B-AS1-related mRNAs.

| ID | Description | GeneRatio | BgRatio | Pvalue | P.adjust | Qvalue | Count |
| --- | --- | --- | --- | --- | --- | --- | --- |
| hsa05340 | Primary immunodeficiency | 16/355 | 38/8081 | 1.27E-12 | 3.41E-10 | 2.99E-10 | 16 |
| hsa04659 | Th17 cell differentiation | 25/355 | 107/8081 | 3.14E-12 | 4.22E-10 | 3.70E-10 | 25 |
| hsa04640 | Hematopoietic cell lineage | 23/355 | 99/8081 | 2.70E-11 | 2.42E-09 | 2.12E-09 | 23 |
| hsa04658 | Th1 and Th2 cell differentiation | 22/355 | 92/8081 | 4.01E-11 | 2.70E-09 | 2.36E-09 | 22 |
| hsa04612 | Antigen processing and presentation | 18/355 | 78/8081 | 4.60E-09 | 2.47E-07 | 2.17E-07 | 18 |
| hsa04650 | Natural killer cell mediated cytotoxicity | 22/355 | 131/8081 | 4.68E-08 | 2.10E-06 | 1.84E-06 | 22 |
| hsa04660 | T cell receptor signaling pathway | 19/355 | 104/8081 | 9.89E-08 | 3.80E-06 | 3.33E-06 | 19 |
| hsa05332 | Graft-versus-host disease | 12/355 | 42/8081 | 1.44E-07 | 4.84E-06 | 4.24E-06 | 12 |
| hsa04940 | Type I diabetes mellitus | 12/355 | 43/8081 | 1.92E-07 | 5.74E-06 | 5.03E-06 | 12 |
| hsa05235 | PD-L1 expression and PD-1 checkpoint pathway in cancer | 17/355 | 89/8081 | 2.42E-07 | 6.50E-06 | 5.70E-06 | 17 |
| hsa05330 | Allograft rejection | 11/355 | 38/8081 | 4.16E-07 | 1.02E-05 | 8.93E-06 | 11 |
| hsa05140 | Leishmaniasis | 15/355 | 77/8081 | 9.57E-07 | 2.14E-05 | 1.88E-05 | 15 |
| hsa04060 | Cytokine-cytokine receptor interaction | 32/355 | 295/8081 | 1.77E-06 | 3.66E-05 | 3.21E-05 | 32 |
| hsa05169 | Epstein-Barr virus infection | 24/355 | 202/8081 | 7.81E-06 | 0.00015 | 0.000132 | 24 |
| hsa05320 | Autoimmune thyroid disease | 11/355 | 53/8081 | 1.46E-05 | 0.000261 | 0.000229 | 11 |
| hsa05321 | Inflammatory bowel disease | 12/355 | 65/8081 | 2.10E-05 | 0.000353 | 0.00031 | 12 |
| hsa03050 | Proteasome | 10/355 | 46/8081 | 2.33E-05 | 0.000369 | 0.000324 | 10 |
| hsa04514 | Cell adhesion molecules | 19/355 | 149/8081 | 2.59E-05 | 0.000388 | 0.00034 | 19 |
| hsa04672 | Intestinal immune network for IgA production | 10/355 | 49/8081 | 4.18E-05 | 0.00059 | 0.000517 | 10 |
| hsa04064 | NF-kappa B signaling pathway | 15/355 | 104/8081 | 4.38E-05 | 0.00059 | 0.000517 | 15 |
| hsa04662 | B cell receptor signaling pathway | 13/355 | 82/8081 | 5.18E-05 | 0.000664 | 0.000582 | 13 |
| hsa04061 | Viral protein interaction with cytokine and cytokine receptor | 14/355 | 100/8081 | 0.000109 | 0.001338 | 0.001173 | 14 |
| hsa05166 | Human T-cell leukemia virus 1 infection | 22/355 | 219/8081 | 0.000236 | 0.002765 | 0.002424 | 22 |
| hsa04062 | Chemokine signaling pathway | 20/355 | 192/8081 | 0.00028 | 0.003139 | 0.002751 | 20 |
| hsa05152 | Tuberculosis | 19/355 | 180/8081 | 0.000335 | 0.003592 | 0.003148 | 19 |
| hsa04145 | Phagosome | 17/355 | 152/8081 | 0.000347 | 0.003592 | 0.003148 | 17 |
| hsa05145 | Toxoplasmosis | 14/355 | 112/8081 | 0.000369 | 0.003676 | 0.003222 | 14 |
| hsa04380 | Osteoclast differentiation | 15/355 | 128/8081 | 0.000466 | 0.004475 | 0.003923 | 15 |
| hsa05323 | Rheumatoid arthritis | 12/355 | 93/8081 | 0.000718 | 0.006664 | 0.005841 | 12 |
| hsa05416 | Viral myocarditis | 9/355 | 60/8081 | 0.001118 | 0.010023 | 0.008785 | 9 |
| hsa05310 | Asthma | 6/355 | 31/8081 | 0.001995 | 0.017308 | 0.015171 | 6 |
| hsa05170 | Human immunodeficiency virus 1 infection | 19/355 | 212/8081 | 0.002419 | 0.020338 | 0.017827 | 19 |
| hsa05162 | Measles | 14/355 | 139/8081 | 0.003084 | 0.024648 | 0.021605 | 14 |
| hsa05150 | Staphylococcus aureus infection | 11/355 | 96/8081 | 0.003115 | 0.024648 | 0.021605 | 11 |
| hsa05020 | Prion disease | 22/355 | 273/8081 | 0.00428 | 0.032898 | 0.028836 | 22 |
| hsa03040 | Spliceosome | 14/355 | 151/8081 | 0.006472 | 0.048364 | 0.042393 | 14 |

**S5 Table.** The KEGG pathway enrichment analysis for AC004687.1-related mRNAs

| ID | Description | GeneRatio | BgRatio | Pvalue | P.adjust | Qvalue | Count |
| --- | --- | --- | --- | --- | --- | --- | --- |
| hsa04640 | Hematopoietic cell lineage | 48/520 | 99/8081 | 1.85E-31 | 5.26E-29 | 4.31E-29 | 48 |
| hsa04060 | Cytokine-cytokine receptor interaction | 77/520 | 295/8081 | 5.39E-28 | 7.65E-26 | 6.27E-26 | 77 |
| hsa05330 | Allograft rejection | 29/520 | 38/8081 | 1.26E-27 | 1.19E-25 | 9.73E-26 | 29 |
| hsa05332 | Graft-versus-host disease | 30/520 | 42/8081 | 4.33E-27 | 3.07E-25 | 2.52E-25 | 30 |
| hsa04672 | Intestinal immune network for IgA production | 31/520 | 49/8081 | 1.91E-25 | 1.09E-23 | 8.90E-24 | 31 |
| hsa04612 | Antigen processing and presentation | 38/520 | 78/8081 | 3.45E-25 | 1.63E-23 | 1.34E-23 | 38 |
| hsa04940 | Type I diabetes mellitus | 27/520 | 43/8081 | 3.56E-22 | 1.45E-20 | 1.18E-20 | 27 |
| hsa04514 | Cell adhesion molecules | 48/520 | 149/8081 | 6.28E-22 | 2.23E-20 | 1.83E-20 | 48 |
| hsa04658 | Th1 and Th2 cell differentiation | 37/520 | 92/8081 | 6.20E-21 | 1.96E-19 | 1.60E-19 | 37 |
| hsa04061 | Viral protein interaction with cytokine and cytokine receptor | 38/520 | 100/8081 | 1.95E-20 | 5.54E-19 | 4.54E-19 | 38 |
| hsa05320 | Autoimmune thyroid disease | 28/520 | 53/8081 | 4.30E-20 | 1.11E-18 | 9.10E-19 | 28 |
| hsa04659 | Th17 cell differentiation | 38/520 | 107/8081 | 3.09E-19 | 7.31E-18 | 5.98E-18 | 38 |
| hsa04062 | Chemokine signaling pathway | 51/520 | 192/8081 | 4.43E-19 | 9.68E-18 | 7.93E-18 | 51 |
| hsa04650 | Natural killer cell mediated cytotoxicity | 41/520 | 131/8081 | 2.35E-18 | 4.77E-17 | 3.91E-17 | 41 |
| hsa05150 | Staphylococcus aureus infection | 34/520 | 96/8081 | 2.74E-17 | 5.20E-16 | 4.26E-16 | 34 |
| hsa05340 | Primary immunodeficiency | 22/520 | 38/8081 | 3.38E-17 | 5.99E-16 | 4.91E-16 | 22 |
| hsa05416 | Viral myocarditis | 27/520 | 60/8081 | 4.20E-17 | 7.02E-16 | 5.75E-16 | 27 |
| hsa05140 | Leishmaniasis | 30/520 | 77/8081 | 9.87E-17 | 1.56E-15 | 1.28E-15 | 30 |
| hsa04380 | Osteoclast differentiation | 38/520 | 128/8081 | 3.14E-16 | 4.69E-15 | 3.84E-15 | 38 |
| hsa05323 | Rheumatoid arthritis | 31/520 | 93/8081 | 5.11E-15 | 7.25E-14 | 5.94E-14 | 31 |
| hsa05321 | Inflammatory bowel disease | 26/520 | 65/8081 | 5.41E-15 | 7.31E-14 | 5.99E-14 | 26 |
| hsa05169 | Epstein-Barr virus infection | 45/520 | 202/8081 | 7.15E-14 | 9.23E-13 | 7.56E-13 | 45 |
| hsa05145 | Toxoplasmosis | 32/520 | 112/8081 | 2.44E-13 | 3.01E-12 | 2.47E-12 | 32 |
| hsa04662 | B cell receptor signaling pathway | 27/520 | 82/8081 | 4.30E-13 | 5.06E-12 | 4.14E-12 | 27 |
| hsa05152 | Tuberculosis | 41/520 | 180/8081 | 4.45E-13 | 5.06E-12 | 4.14E-12 | 41 |
| hsa04145 | Phagosome | 36/520 | 152/8081 | 3.70E-12 | 4.04E-11 | 3.31E-11 | 36 |
| hsa04064 | NF-kappa B signaling pathway | 29/520 | 104/8081 | 6.51E-12 | 6.85E-11 | 5.61E-11 | 29 |
| hsa05310 | Asthma | 16/520 | 31/8081 | 8.42E-12 | 8.54E-11 | 6.99E-11 | 16 |
| hsa04660 | T cell receptor signaling pathway | 28/520 | 104/8081 | 3.82E-11 | 3.74E-10 | 3.06E-10 | 28 |
| hsa05166 | Human T-cell leukemia virus 1 infection | 41/520 | 219/8081 | 3.33E-10 | 3.15E-09 | 2.58E-09 | 41 |
| hsa05144 | Malaria | 17/520 | 50/8081 | 5.78E-09 | 5.30E-08 | 4.34E-08 | 17 |
| hsa05322 | Systemic lupus erythematosus | 29/520 | 136/8081 | 6.65E-09 | 5.90E-08 | 4.84E-08 | 29 |
| hsa04630 | JAK-STAT signaling pathway | 32/520 | 162/8081 | 8.00E-09 | 6.89E-08 | 5.64E-08 | 32 |
| hsa05170 | Human immunodeficiency virus 1 infection | 34/520 | 212/8081 | 5.67E-07 | 4.74E-06 | 3.88E-06 | 34 |
| hsa05133 | Pertussis | 18/520 | 76/8081 | 1.01E-06 | 8.17E-06 | 6.69E-06 | 18 |
| hsa05142 | Chagas disease | 21/520 | 102/8081 | 1.53E-06 | 1.21E-05 | 9.88E-06 | 21 |
| hsa05235 | PD-L1 expression and PD-1 checkpoint pathway in cancer | 19/520 | 89/8081 | 2.73E-06 | 2.09E-05 | 1.71E-05 | 19 |
| hsa04610 | Complement and coagulation cascades | 18/520 | 85/8081 | 5.63E-06 | 4.21E-05 | 3.44E-05 | 18 |
| hsa05164 | Influenza A | 27/520 | 171/8081 | 1.13E-05 | 8.24E-05 | 6.75E-05 | 27 |
| hsa05143 | African trypanosomiasis | 11/520 | 37/8081 | 1.30E-05 | 9.21E-05 | 7.55E-05 | 11 |
| hsa05135 | Yersinia infection | 22/520 | 137/8081 | 5.63E-05 | 0.00039 | 0.000319 | 22 |
| hsa04670 | Leukocyte transendothelial migration | 19/520 | 114/8081 | 0.000108 | 0.000727 | 0.000596 | 19 |
| hsa05167 | Kaposi sarcoma-associated herpesvirus infection | 26/520 | 193/8081 | 0.000251 | 0.001659 | 0.001359 | 26 |
| hsa04621 | NOD-like receptor signaling pathway | 24/520 | 181/8081 | 0.000545 | 0.00342 | 0.002802 | 24 |
| hsa05162 | Measles | 20/520 | 139/8081 | 0.000545 | 0.00342 | 0.002802 | 20 |
| hsa05163 | Human cytomegalovirus infection | 28/520 | 225/8081 | 0.000554 | 0.00342 | 0.002802 | 28 |
| hsa04668 | TNF signaling pathway | 17/520 | 112/8081 | 0.000755 | 0.004562 | 0.003737 | 17 |
| hsa04611 | Platelet activation | 18/520 | 124/8081 | 0.000916 | 0.005237 | 0.00429 | 18 |
| hsa04620 | Toll-like receptor signaling pathway | 16/520 | 104/8081 | 0.000922 | 0.005237 | 0.00429 | 16 |
| hsa04625 | C-type lectin receptor signaling pathway | 16/520 | 104/8081 | 0.000922 | 0.005237 | 0.00429 | 16 |
| hsa05134 | Legionellosis | 10/520 | 57/8081 | 0.003099 | 0.017257 | 0.014136 | 10 |
| hsa05202 | Transcriptional misregulation in cancer | 22/520 | 192/8081 | 0.005816 | 0.031765 | 0.02602 | 22 |
| hsa04210 | Apoptosis | 17/520 | 136/8081 | 0.006202 | 0.033235 | 0.027223 | 17 |
| hsa04666 | Fc gamma R-mediated phagocytosis | 13/520 | 97/8081 | 0.00894 | 0.047017 | 0.038513 | 13 |

**S6 Table.** The KEGG pathway enrichment analysis for PCED1B-AS1-related mRNA.
